# AI-discovered tuning laws explain neuronal population code geometry

**DOI:** 10.1101/2025.11.12.688086

**Authors:** Reilly Tilbury, Dabin Kwon, Ali Haydaroğlu, Jacob Ratliff, Valentin Schmutz, Matteo Carandini, Kevin Miller, Kim Stachenfeld, Kenneth D. Harris

## Abstract

The activity of visual cortical neurons forms a population code representing image stimuli. There is, however, a discrepancy between our understanding of this code at the single-cell and population levels: direct measurements indicate the population code is high-dimensional, but established models of single-cell tuning give rise to low-dimensional codes. We reconciled this discrepancy by developing an AI science system to find a new parsimonious, interpretable equation for visual cortical orientation tuning. Candidate equations were expressed as short computer programs and evolved by Large Language Models (LLMs) using graphical diagnostics. The resulting equation not only improved single-cell fits, but also accurately modelled the population code’s high-dimensional geometry. A novel parameter of the AI-discovered equation, which controls single-cell tuning smoothness, gives rise to high-dimensional population codes. The same parameter drives high-dimensional coding in head-direction cells, suggesting a common coding strategy across brain regions. We used this equation to hypothesize a circuit mechanism generating high-dimensional population codes, and to demonstrate the advantages of these codes in a simulated hyperacuity task. These results show that tuning smoothness has a key role in controlling population code geometry, and demonstrate how AI equation discovery can deliver interpretable models accelerating scientific understanding in neuroscience and beyond.

---

The visual cortex comprises myriad neurons, whose joint activity forms a “population code” representing image stimuli. The properties of the cortical population code determine the types of computations that can be performed by downstream brain structures. Theoretical work has long recognized the advantages of high-dimensional codes, which allow downstream neurons to flexibly read out diverse features by simple linear integration ^1–7^. The cortex can form such high-dimensional codes by embedding a manifold of possible sensory stimuli nonlinearly into a high-dimensional space of neuronal activity patterns. Nonlinear transformation of sensory inputs into high-dimensional codes, which allow linear readout of behaviorally-relevant features, has been proposed as a primary function of the cerebral cortex ^7–14^.

The dimensionality of the neural code for a stimulus set can be revealed by principal component analysis (PCA) of neuronal response vectors. The PCA eigenspectrum provides a sequence of numbers *V*_*n*_ measuring the variance of activity along the *n*^*th*^ embedding dimension of the response manifold, whose dependence on *n* characterizes the population code’s dimensionality. If the eigenspectrum decays exponentially, e.g. *V*_n_ ∼*e*^−*n*/*M*^ or 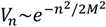 this indicates a low-dimensional code, with *M* setting a dimension scale beyond which variances rapidly vanish. If instead the eigenspectrum decays as a power law *V*_*n*_∼*n*^−*α*^, this indicates a higher-dimensional code with no single characteristic dimension, as has been observed in visual cortex ^15^.

The population code emerges from the tuning of individual neurons, but there is a discrepancy between our understanding of sensory tuning and population coding in visual cortex. This discrepancy is most evident for responses to moving grating stimuli of varying orientation and direction. Single-neuron orientation tuning has been measured many times since the 1960s, typically by varying direction of motion *θ* in steps of 15° or 30°, and fitting the results with a smooth function, such as the “double Gaussian”^16,17^:

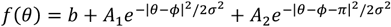

This equation predicts a low-dimensional population code, but the actual code is high-dimensional. To expose this discrepancy, we analyzed 2-photon calcium recordings of visual cortical responses to drifting gratings of fine angular spacing (Fig. 1a,b). To probe the high-dimensional structure of this data, we developed a high-precision regularized, cross-validated variant of PCA (rcvPCA; Fig. S1; Methods), which estimates the eigenspectrum of stimulus responses discarding stimulus-independent variability. This revealed a power-law eigenspectrum decay with exponent ≈ 4 (Fig. 1c). Next, we fit double-Gaussian tuning curves to the responses of each neuron (Fig. 1b) and computed the PCA eigenspectrum of the resulting population of tuning curves, obtaining exponential decay indicative of a low-dimensional code (Fig. 1c,d). This discrepancy could be visualized by projecting population activity and its double-Gaussian fit onto principal component axes (Fig. 1e,f). Actual population activity shows clear, repeatable structure in higher dimensions (e.g. dimension 24, Fig. 1e), which the double-Gaussian equation fails to reproduce (Fig. 1f). This discrepancy is not restricted to Gaussian tuning: mathematically, no family of tuning curves which is infinitely differentiable can produce eigenspectra with power-law decay (Supplementary Text 1a,b; Ref. ^10^). Because current orientation tuning models have this property, explaining high-dimensional population coding requires a new equation for single-cell tuning curves.

**Figure 1.**
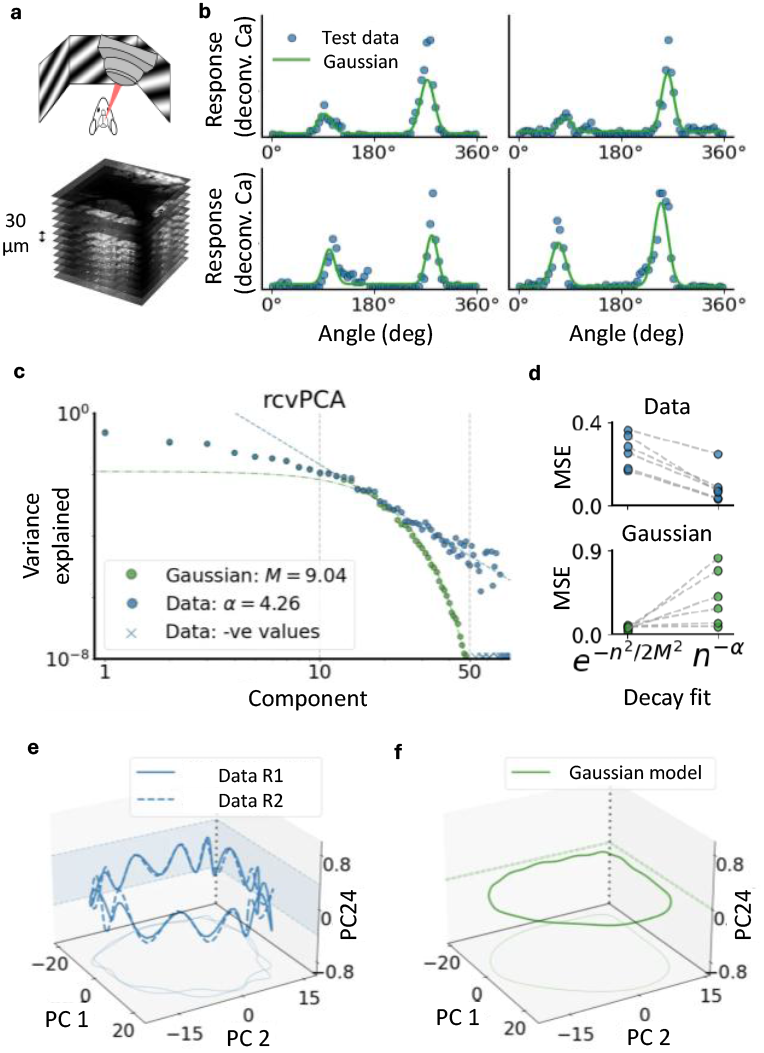
Oriented stimuli produce a high-dimensional population code not captured by standard tuning curve models. **a**, Visual cortical population activity was recorded in awake mice passively presented with full-field drifting grating stimuli. **b**, Responses of four example cells as a function of grating direction (blue dots) together with double Gaussian model fits (green curves). **c**, Population eigenspectrum for an example experiment (blue dots; blue crosses indicate dimensions for which rcvPCA estimates are negative) and eigenspectrum of a population of double-Gaussian tuning curves fit to the recorded cells (green dots). Blue dashed line: *n*^−*α*^ fit to actual eigenspectrum; green dashed line: 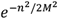 fit to eigenspectrum of Gaussian tuning curves, both with parameters estimated in the region 10 < n < 50. **d**, Mean-square error of 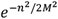 and *n*^−*α*^ fits to eigenspectra of actual data (upper; each pair of blue points joined by dashed line is one recording) and Gaussian model (lower). **e**, Population responses projected onto principal components 1, 2, and 24. Two independent repeats of experimental data (R1 and R2) show consistent high-dimensional structure. **f**, Same as (e), for the population of Gaussian model fits, which shows low-dimensional structure.

We had not found a suitable equation with traditional means, so turned to a new approach, which we term AI equation discovery (Fig. 2a). This approach differs from previous uses of AI in studying neural tuning, which involve training deep networks to match visual responses ^18–26^, resulting in networks which can reconstruct responses with high accuracy, but are challenging to interpret. AI equation discovery instead aims to produce a parsimonious, interpretable equation summarizing how neurons respond to stimuli. In contrast to symbolic regression and other approaches which evolve equations or programs as syntax trees subject to mutation and cross-over ^27–31^, this approach takes advantage of the capacity of modern LLMs to reason scientifically and interpret graphical representations of data ^32–39^.

**Figure 2.**
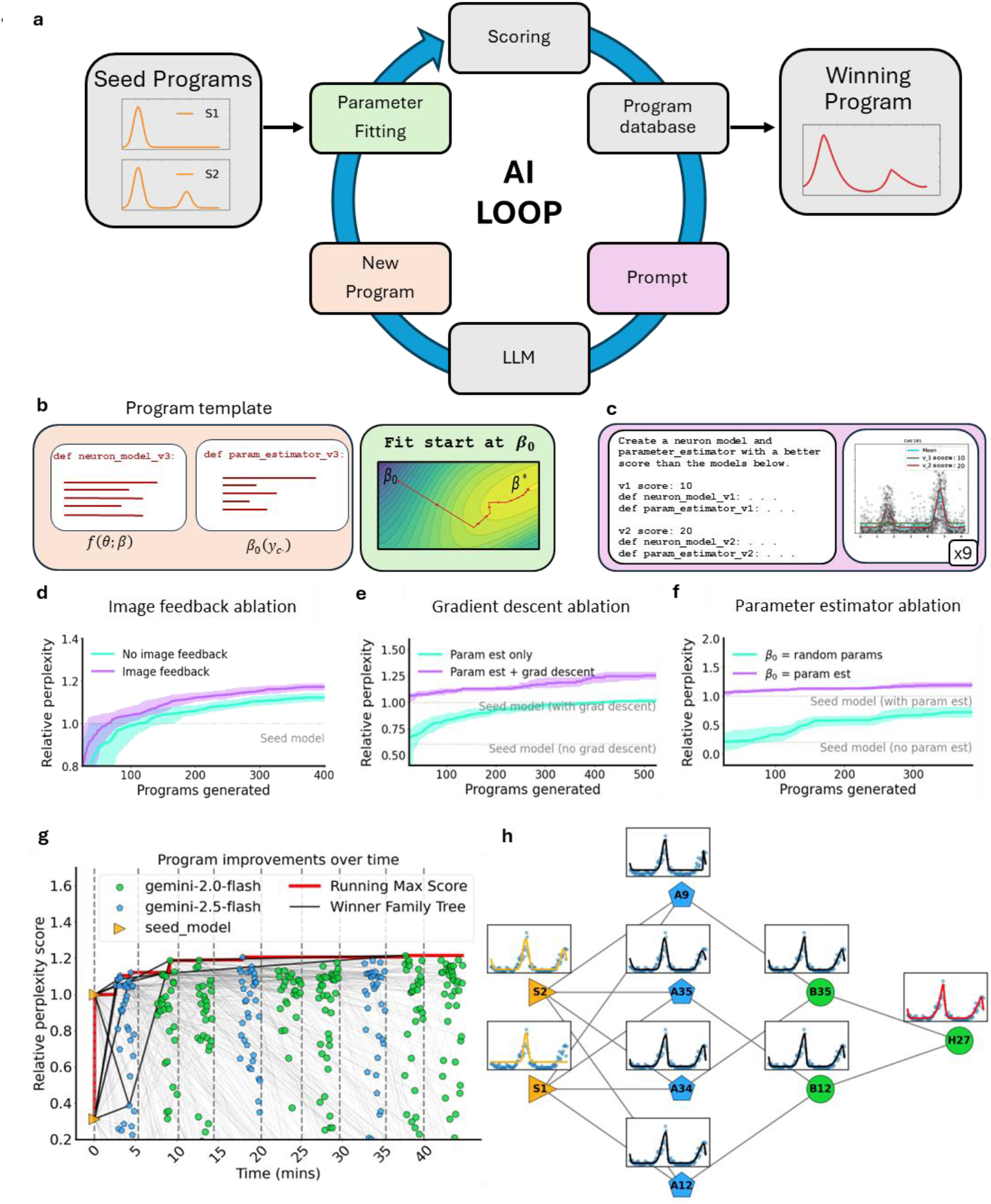
AI equation discovery system. **a**, System flowchart. Tuning curve equations are represented as Python programs. An LLM is repeatedly prompted to improve on two currently high-scoring programs, and the result is scored by complexity-penalized cross-validated squared error, and entered into a database (“AI LOOP”). Finally, a winner is selected using held-out cells. **b**, Each program comprises a tuning function *f*(θ, *β*) predicting the response of a neuron with tuning parameters *β* to stimulus θ, and a parameter estimator 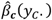 that provides a starting point for gradient optimization of each cell’s tuning parameter *β*_*c*_ given its measured responses *y*_*C*_ . **c**, Prompts quote two high-scoring programs and their scores, and contain graphical diagnostics comparing these programs’ predictions to actual responses for nine randomly selected neurons. **d**, Ablation test comparing text-only feedback (cyan) vs. system including graphical diagnostics (magenta). **e**, Ablation test comparing system without gradient descent (cyan) to full system (magenta). **f**, Ablation test comparing system with random search initialization (cyan) to that with the parameter estimator (magenta). Solid lines in D-F show median fitness of the top cohorts (defined as the top 24 programs of each condition, pooled across 4 independent evolutions), error bars show 25^th^ and 75^th^ percentiles of the top cohorts. **g**, Summary of programs generated during equation discovery. X-axis: wall-clock time of program generation. Y-axis: perplexity (i.e. exponential of negative error), relative to that of double-Gaussian seed model. Red line: running-best score. Black lines: winner’s family tree. **h**, Family tree of the winning model (right), and fits of each model in it for an example cell. For the equations of these models, see Supplementary Text 3.

We implemented this using an evolutionary approach ^40–43^ (Fig. 2a). Equations for tuning curves are expressed as Python programs predicting a cell’s firing rate response to each stimulus, given cell-specific tuning parameters. The programs are scored by cross-validated mean-squared error, penalized by program complexity (Fig. S2), and all programs and scores are entered into a database. Programs are evolved by repeatedly sampling two high-scoring “parent” programs from the database, and prompting an LLM to generate a new, higher-scoring “child” model. The prompt quotes the code of both parent programs, as well as graphical diagnostics comparing the parent models to experimental data (Figs. 2b,c, S3), a new feature which we hypothesized might help find better solutions than scores alone. The system ran for approximately 45 minutes of wall-clock time, wherein 576 programs were created and evaluated at a cost of $8.25 in LLM tokens, substantially less than previous approaches requiring up to millions of program evaluations ^40–43^. Ablation tests indicated significant improvements from giving the LLMs graphical model diagnostics, as well as another new feature whereby the LLMs also write code to estimate an approximate parameter fit, used as a gradient descent starting point to avoid local minima (Figs. 2d-f, S4). For example prompts and results, see Supplementary Text 2.

The winning model was identified as that with best cross-validated performance on a set of test cells held out from model evolution. We ran the system 8 times, of which 6 runs converged on the same equation

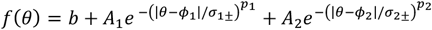

This has three key differences to the original double-Gaussian model, which were progressively added over several generations (Figs. 2g,h; Supplementary Text 3). Most importantly, the power of 2 in the Gaussian has been replaced by parameters *p*_1_ and *p*_2_, which control the shape of the two peaks (Fig. 3a). If *p* = 2, this yields the usual infinitely differentiable Gaussian function. When *p* < 2, the curve is “peaky”, and not infinitely-differentiable; when *p* < 1 it has cusp-shaped peaks, allowing it to fit large responses to exactly preferred stimuli which appeared to be outliers when fit by the Gaussian (Fig. 3b-d), but occurred specifically at each cell’s preferred orientation, indicating a reliable feature of neuronal tuning (Figs. S5,S6). Second, the model allows the two preferred directions *ϕ*_1_ and *ϕ*_2_ to take arbitrary values rather than be 180° apart as in the original model. Finally, *σ*_1±_ and *σ*_2±_ represent two parameters each (*σ*_1±_ = *σ*_1−_ when *θ* < *ϕ*_1_ and *σ*_1±_ = *σ*_1+_when *θ* > *ϕ*_1_), allowing different widths to the left and right of the tuning peak, a phenomenon first noticed in the 1970s (Refs. ^44,45^) but rarely discussed since. The parameters varied widely between cells (Fig. S7), and ablation tests showed that all three features improved cross-validated explained variance (Fig. 3e), although this improvement was relatively modest as the new parameters improved the fit of only stimuli close to the tuning-curve peaks.

**Figure 3.**
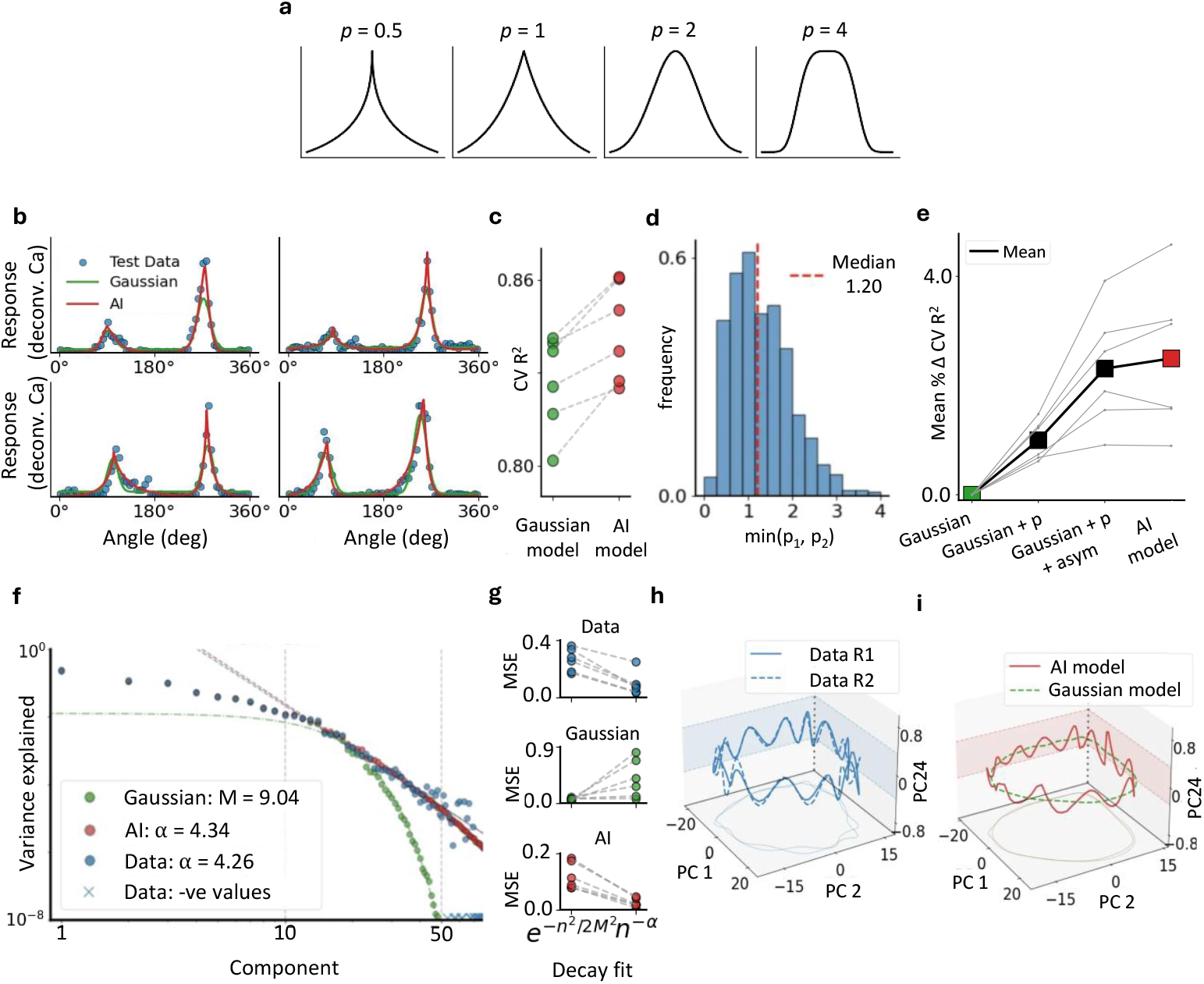
AI-discovered tuning function explains high dimensional population code. **a**, Shape of function peaks: cusp for p=0.5, triangular for p=1, smooth for p=2 and flat for p=4. **b**, AI-derived tuning function 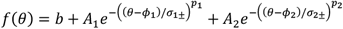 fit to the same example cells as in Fig. 1B. Blue points: experimental data; red curve: AI model; green curve: double-Gaussian model. **c**, Fit quality of double-Gaussian and AI models, as assessed by cross-validated *R*^2^. Circles: individual experiments. **d**, Histogram over all cells of exponent *p* (minimum over the two peaks, which determines the power-law exponent). **e**, Quantifying each parameter’s effect by systematically ablating parameters from the equation. Thin curves show change in explained variance over double-Gaussian model, averaged over cells for single experiments; thick curve: mean. **f**, PCA eigenspectrum of AI model (red points) with eigenspectra of actual data (blue) and Gaussian model (green). **g**, mean-square error of 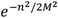 and *n*^−*α*^ fits to each spectrum, as in Figure 1D. **h**,**i**, Population responses and AI model projected onto principal components 1, 2, and 24, as in Fig. 1e,f.

The new AI-derived tuning equation, unlike the traditional model, predicted a high-dimensional population code. To show this, we fit AI-derived tuning curves to each cell. The PCA eigenspectrum of the population of AI model tuning curves closely matched the *n*^−4^ powerlaw structure of the original data (Fig. 3f,g) and higher dimensionality could be visualized by principal component projection (Fig. 3h,i). Because the AI system found an explicit equation for the tuning curves, we could explain this result mathematically, proving that for *p* < 2, tuning curves of the form 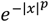 generate population codes with power-law eigenspectrum decay with exponent 2(*p* + 1), due to their non-infinitely-differentiable peaks (Supplementary Text 1b); for double-peaked tuning curves, the exponent is set by the smaller *p*. The values of *p* fit to V1 data were centered around 1 (Fig. 3d), and thus consistent with the observed eigenspectrum power-law exponent of around 4. Notably, this dramatic improvement in predicting the high-dimensional structure of the population code occurred even though the LLM’s objective function was based on single-cell error, not population geometry, and even though the AI system produced only a relatively modest improvement in its single-cell-based objective function.

To test whether peaky tuning might generate high-dimensional population codes in brain systems beyond visual cortex, we analyzed data from head-direction cells (Fig. 4). Head-direction cells also encode an angular variable, the head direction of a subject physically moving through space. We analyzed data collected from head-direction cells in the thalamus and post-subiculum of mice freely exploring an open field environment ^46^, fitting each recorded cell with the AI-derived tuning equation (modified to have only one peak, matching the single-peaked nature of head-direction tuning curves). Individual cells’ tuning curves were better fit by the AI model than Gaussian tuning (Fig. 4a,b). Applying rcvPCA to a population of head direction cells (pooling cells across mice to more accurately estimate higher dimensions) revealed a high-dimensional power-law eigenspectrum, that was again accurately captured by the AI-derived tuning equation (Fig. 4c,d).

**Figure 4.**
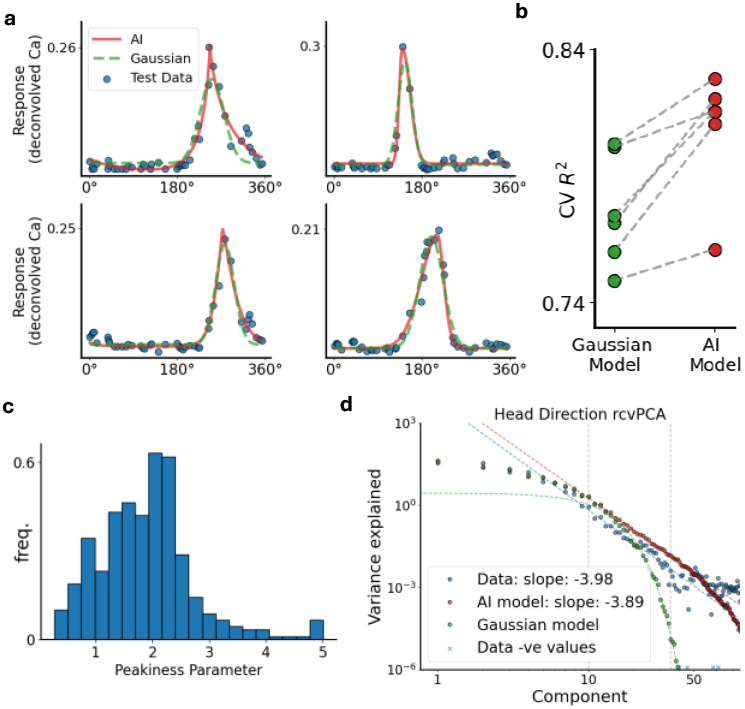
Analysis of head-direction cell data. **a**, Tuning of four example head direction cells (blue points), together with AI (red) and Gaussian (dashed green) tuning curves. **b**, Fit quality of single-Gaussian and single-peaked AI model, as assessed by cross-validated *R*^2^. Circles: individual mice. **c**, Histogram of *p* parameter across cells. **d**, Eigenspectrum of HD cell population activity (blue) together with Gaussian (green) and AI (red) models. Blue and Red dashed lines: *n*^−*α*^ fit to population eigenspectrum and AI model; green dashed line: 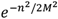 fit to Gaussian model.

To investigate the computational advantages of this high-dimensional population code for angular data, we took advantage of the explicit equation discovered by the AI system. We modeled a hyperacuity task, where a simulated agent is required to discriminate whether the stimulus orientation is in a region smaller than the tuning curve half-width, which can be performed by humans and animals ^47–49^. We considered a population of symmetrical tuning curves of the form 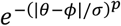, with fixed half-width *σ* =15° and evenly-distributed tuning curve centers, and asked how well a linear decoder of this activity could discriminate orientations closer or further than 3° from a target, for different values of *p* (Fig. 5a). We simulated a downstream readout that learns by Hebbian plasticity and found that it showed better performance and faster learning for sharp tuning curves with *p*=0.5 or 1.0 than for traditional smooth tuning curves with *p*=2.0 (Fig. 5b).

**Figure 5.**
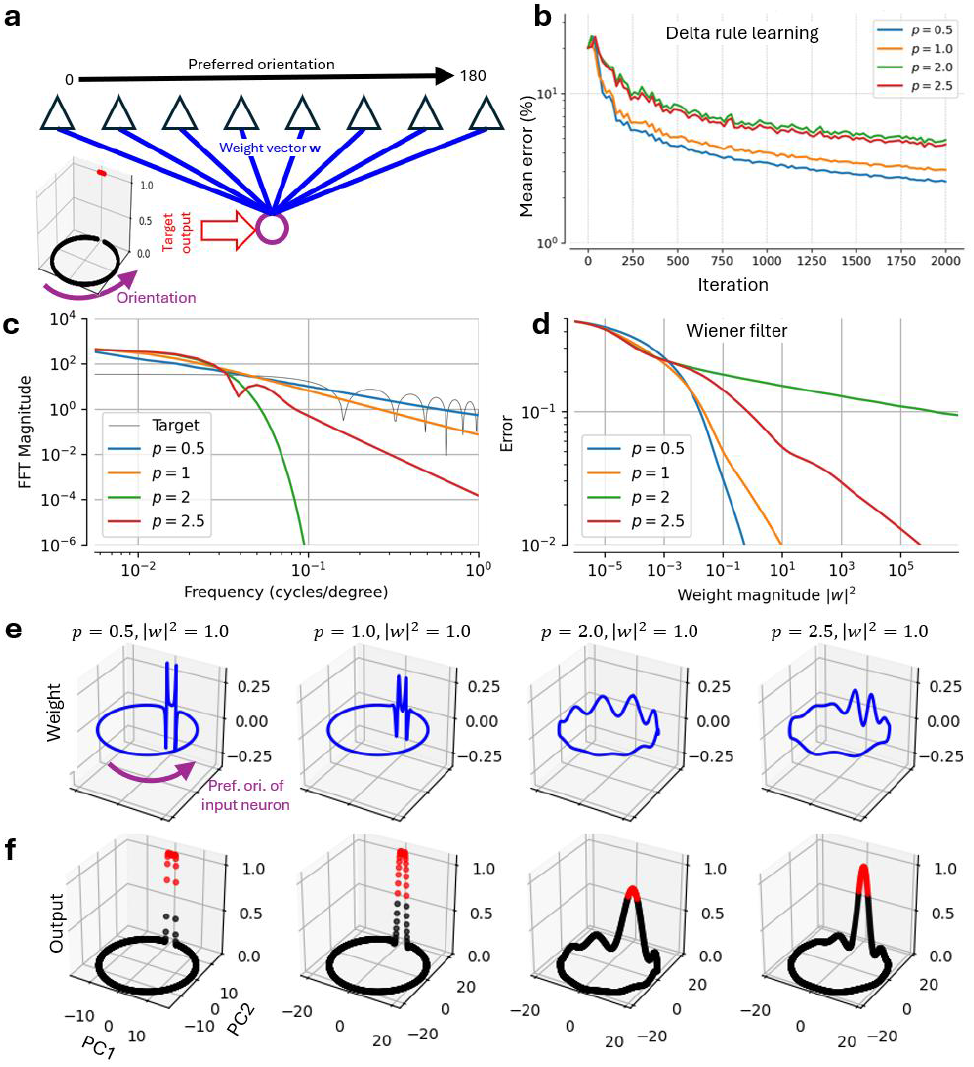
Effect of tuning peakiness on a simulated hyperacuity task. **a**, Simulation set up: a population of tuning curves of the form 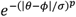 with evenly distributed preferred orientations, width *σ* =15°, and varying *p*, are linearly read out to discriminate a target function. The target function (inset) is 1 for orientations within 3° of a target angle (red points) and 0 otherwise (black points). **b**, Readout error vs. time for a Hebbian learning rule, for different values of p. **c**, Fourier transform of the target function (black) and of tuning curves for different values of *p* (colors). **d**, Mean squared error achievable with optimal weight choice, as a function of weight norm (|*w*|^2^) for different values of p. **e**, Optimal readout weights with fixed norm (|*w*|^2^ = 1), for different values of *p*. X- and y-axes: sine and cosine of preferred angle *ϕ* of source neuron. Z-axis: weight value. **f**, Optimal predictions using the weights shown in (e). X- and y-axes: first two principal components of population activity, reflecting sine and cosine of stimulus angle (note total variance for these components when p=0.5). Z-axis: optimal predictor of boxcar target. Black points: angles for which target is 0, red points, angles for which target is 1.

Thanks to the AI-derived equation, these computational advantages of sharp tuning curves can be explained analytically. Because the neurons are arranged on a circle of preferred orientation *ϕ*, a weighted sum of their neural activity is the circular convolution of the tuning curve and the weight function, which in the Fourier domain becomes a multiplication: *F*[∫ *w*(*ϕ*)*f*(*θ* − *ϕ*)*dϕ*] = *F*[*w*]*F*[*f*]. Accurate readout with reasonably-sized weights therefore requires the Fourier transform of the tuning curve to be comparable in magnitude to that of the target function. The boxcar target function has Fourier decay ∼*n*^−1^, while the modeled tuning curves decay as *n*^−(*p*+1)^ for *p* ≠ 2, but exponentially fast for *p* = 2 (Fig. 5c). Thus, accurate reconstruction of the target function with reasonable weights is possible for small values of *p*, but not for *p* = 2 (Fig. 5d). To illustrate this, we found the exact optimal solutions for fixed weight vector norm using a Wiener filter, and found that for *p*=0.5 or 1, accurate solutions are obtained by integrating the activity of neurons with preferred orientation near the target value (Fig. 5e,f). For *p*=2 however, the optimal weights are a widespread and oscillatory function of preferred orientation, and reconstruction is substantially worse. Intriguingly, even the case of *p*=2.5, whose tuning curves are flatter than Gaussian but not infinitely differentiable, yielded better performance than the smooth Gaussian with *p*=2. Thus, the Gaussian tuning curve is exceptionally unsuited for making fine orientation discriminations such as required for hyperacuity, while peaky tuning curves allow better performance.

Finally, we considered what circuit mechanisms might generate peaky stimulus tuning, even in cases where external inputs are infinitely differentiable (Fig. 6). A single-cell nonlinearity cannot generate this peaky tuning robustly. Indeed, by Taylor’s theorem, an infinitely-differentiable input tuning curve *f*(*θ*) can be approximated near its maximum *θ*_0_ by a quadratic *f*(*θ*) ≈ *A*(1 − *β*^2^(*θ* − *θ*_0_)^2^). Transforming this to *A*(1 − *β*|*θ* − *θ*_0_|), which more closely matches the observed tuning curves, requires a transformation 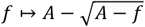, which has infinite slope at *f* = *A* and is undefined for *f* > *A*. There are no known single-cell processes that generate this kind of a transformation, especially because the maximum input *A* would not be constant: it would vary due to changes in input strength. Nevertheless, we illustrate how such a transformation could be performed by a recurrent network, using an exactly solvable variant of the “stabilized supralinear network” model ^50^ (Fig. 6a-d). The model contains excitatory populations tuned to each stimulus, connected by supralinear recurrent excitation which would alone cause activity to explode to infinity, but is checked by high-gain feedback inhibition. We proved analytically that this model implements transformation of the form 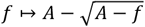, robustly for different values of the maximum input *A* (Supplementary Text 1c) and confirmed in simulation that it transforms Gaussian to peaky tuning curves (Fig. 6e,f). Furthermore, the model predicts narrowing of tuning with time (Fig. 6e), which has been observed in visual and also auditory cortex ^51,52^.

**Figure 6.**
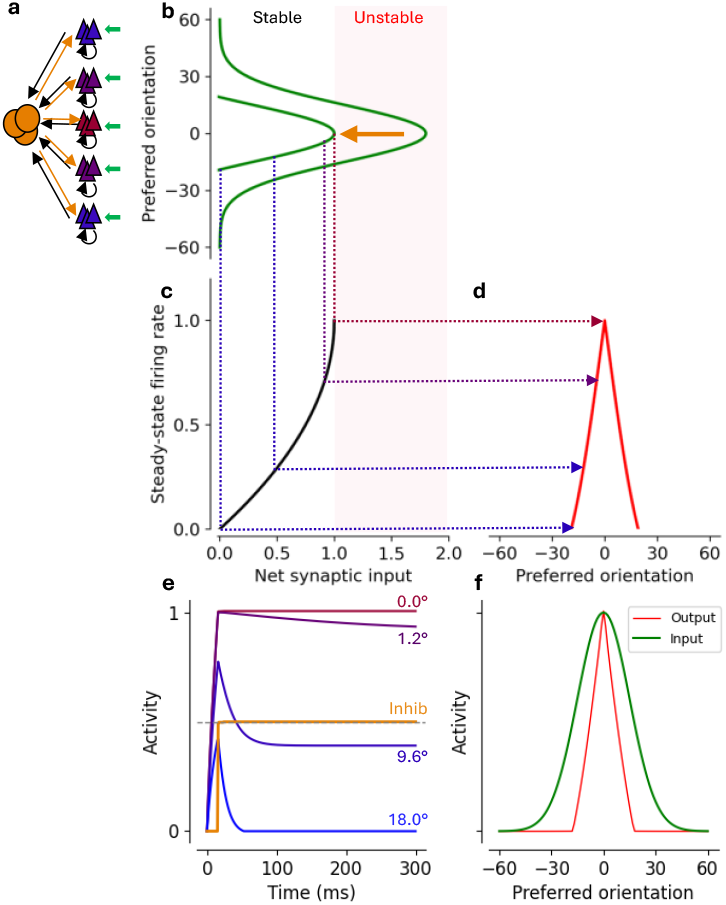
Stabilized supralinear network model transforms smooth inputs to peaky outputs. **a**, Hypothesized circuit mechanism. Excitatory neurons receiving similarly-tuned inputs (magenta to blue) are supralinearly recurrently connected, with activity stabilized by global feedback inhibition (orange). **b**, Synaptic input (x-axis) as a function of excitatory neurons’ preferred orientation (y-axis) for a stimulus of 0°; note that in this plot, x depends on y. Rightmost green curve: external excitation; leftmost green curve: net synaptic input after feedback inhibition (orange arrow). **c**, In the exactly solvable model, supralinear recurrent excitation transforms a net synaptic input *x* into a steady-state firing rate of 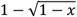 (black curve). For net input > 1, recurrent excitation is unstable, but feedback inhibition causes net input to the most excited cells to settle just below the threshold of instability. **d**, Predicted steady state firing rate in the exactly solvable model given a 0° stimulus, as a function of neuronal preferred orientation. Dashed lines connecting (b), (c), and (d) indicate transformation of Gaussian-tuned inputs to peaky outputs, with colors reflected preferrerd orientation as in (a). **e**, Simulation results. Magenta-to-blue lines: firing rate of excitatory populations receiving inputs of preferred orientation indicated by colors as in (a). Orange line: inhibitory firing rate. **f**, Simulated tuning curves for external input (green) and steady-state firing rate (red).

In summary, we used an AI-science system to discover an equation for direction tuning in mouse visual cortex. The AI system discovered an equation for “peaky” tuning curves, whose non-infinite-differentiability produced a high-dimensional population code, in contrast to the low-dimensional code produced by standard Gaussian tuning. Peaky tuning curves also underpin high-dimensionality in the mouse head-direction system, and future research can investigate whether they are found in a wider range of brain circuits. We illustrated how the high-dimensional codes produced by peaky tuning could allow flexible readout in a hyperacuity task and constructed an exactly solvable circuit model for producing them based on stabilized supralinear excitation. These results suggest that the smoothness of neuronal tuning curves has an important role in controlling the dimensionality of the population code, and thus for understanding how cortical circuits process information.

These results also illustrate one way that AI reasoning can now contribute to scientific research. Fitting simple equations to experimental data can rapidly advance scientific understanding, and the equation found by AI in this study provided an analytical understanding we would not have gained from modeling tuning curves with artificial neural networks, however accurate these models had been. The equation the AI system found is simple, and, in principle, the humans on our team could have come up with it unaided. Nevertheless, we only found it after constructing the AI system. Finding equations summarizing experimental data is a task well-suited to AI science, because the performance of candidate equations can be numerically quantified. We suggest that AI agents will provide a powerful addition to human research teams in cases like this, in neuroscience and beyond.

## Acknowledgements

This work was funded by UKRI (Frontier Award EP/X022366/1 to MC), BBSRC (grant BB/W019884/1 to MC), the Wellcome Trust (Investigator Award 223144/Z/21/Z to MC and KDH), and the ERC (101097874 to KDH). AH is supported by a studentship from the Gatsby Charitable Foundation (GAT3755) and the Wellcome Trust (219627/Z/19/Z). VS is supported by a Royal Society Newton International Fellowship (NIF\R1\231927) and a fellowship from the Swiss National Science Foundation (grant no. 222150).

## Author contributions

Conceptualization: RT, AH, VS, MC, KM, KS, KDH; Resources: MC, KDH; Data curation: RT, DK, AH, JR; Software: RT, DK, KDH; Formal analysis: RT, KDH; Supervision: MC, KM, KS, KDH; Funding acquisition: MC, KDH; Validation: RT; Investigation: RT, DK, AH, JR, KDH; Visualization: RT, DK, KDH; Methodology: RT, KDH; Writing – original draft: RT, KDH; Project administration: KDH; Writing – review and editing: all authors.

## Data and materials availability

All data analyzed in this study are available online at https://figshare.com/s/be52c5b3bb6577fe768e. Code is available at https://github.com/reillytilbury/EDGAR and https://github.com/kdharris101/peaky-tuning-models.

## Methods

### Experimental Methods and datasets analyzed

All experimental procedures were conducted according to the UK Animals Scientific Procedures Act (1986). Experiments were performed at University College London under personal and project licenses released by the Home Office following appropriate ethics review.

The data analyzed here came from four types of experiment. The first three are subtly different types of 2-photon calcium imaging of drifting grating data. The AI system discovered the tuning equation using only dataset 1, and datasets 2 and 3 were used to validate its generality. The fourth is head-direction cell recordings; AI equation discovery was not rerun on this data, instead the equation derived from dataset 1 was tested, after modification to contain only 1 peak.

#### Dataset 1

Dataset 1 was collected and shared by Stringer et al. ^53^, wherein the methods are fully described, and was downloaded from https://janelia.figshare.com/articles/dataset/Recordings_of_20_000_neurons_from_V1_in_response_to_oriented_stimuli/8279387. This data contains activity of visual cortical excitatory neurons, recorded using a custom 2-photon mesoscope. Gratings of spatial frequency 0.05 cycles/degree were presented for 750 ms, at random floating-point directions between 0° and 360° on each trial.

For analysis, the floating-point direction angles were binned into 256 evenly-spaced direction bins, so that each bin contained at least 6 different trials. If more than 6 trials were present for a direction bin, stimuli were randomly discarded to make the remaining number of stimuli divisible by 6, and then averaged in random combination to make exactly 6 repeats per bin.

#### Dataset 2

Dataset 2 was collected in-house. We used a ThorLabs Bergamo 2 microscope to record activity in 2 mice, presenting full-field drifting sinusoidal gratings of 1024 directions evenly-sampling the 360° possible. The stimuli were shown in 3 blocks of exactly 1024 evenly-spaced orientations, in a different random order each time.

Transgenic adult mice expressing TdTomato in inhibitory neurons (Gad-Cre {Jax #010802} x Ai9 {Jax # 007909}) underwent surgery in a stereotaxic frame under isoflurane anaesthesia (5% for induction, 1 to 1.5% maintenance), using previously-described methods ^15,54^. Briefly, a 3 mm craniotomy was made in the skull over primary visual cortex using a biopsy punch. To express GCaMP6s, AAV1-Syn-GCaMP6s (Addgene #100843) was injected intracerebrally through the open craniotomy using a Nanoject II and sharpened glass pipette while the brain surface was covered with hydrated surgical gelatin sponge. A set of glass coverslips was implanted over the craniotomy, attached with veterinary cyanoacrylate glue and secured to the skull with Metabond dental cement alongside a metal headbar ^55^. Animals were allowed to recover for at least one week before habituation to head fixation and imaging.

On the imaging day, animals were head fixed under a ThorLabs Bergamo 2 microscope and allowed to run freely on a cylindrical treadmill. A panel of 1024 evenly spaced, sinusoidal drifting gratings spanning 360° was presented on a 3-monitor array covering a majority of the visual field. The stimuli were shown in 3 blocks of 1024 stimuli in a different random order each time. Gratings had a spatial frequency of 0.05 cycles per visual degree, a temporal frequency of 2 cycles per second, and 100% contrast. Stimuli were presented for 1 second with an intertrial delay of 1 second. Images were acquired in 12 planes spaced 30 microns apart vertically and sampled at a volume rate of 2 Hz. Data were processed using Suite2p ^56^ and all ROIs marked as cells were taken for analysis. Responses were taken as the average deconvolved calcium signal during the duration of visual stimulation.

For analysis, stimuli were binned into 256 evenly spaced direction bins, resulting in 4 neighboring stimulus directions per bin per repeat. The responses of the two odd and even directions in each bin were averaged, resulting in 6 repeats per bin.

#### Dataset 3

Dataset 3 we used a custom light-beads microscope ^57^ to record responses to 2 blocks of repeats full-field drifting square grating stimuli, of random integer directions ranging from 0° to 359°, in one mouse. This data has been previously analyzed in Ref. ^58^, wherein the methods are described, and are available at https://figshare.com/s/aac82eac1829b7ec406e.

These data were binned similarly to Dataset 1, but with 90 bins, again resulting in exactly 6 repeats per bin.

#### Dataset 4

The fourth dataset was the activity of head-direction cells in freely-moving exploring mice, collected and shared by Peyrache et al. ^46^, and was downloaded from https://crcns.org/data-sets/thalamus/th-1. The dataset comprises 42 recording sessions obtained from seven freely moving mice while they foraged for food in an open arena (53 × 46 cm). Of these, we focused on six mice (12, 17, 20, 24, 25, and 28), comprising 39 sessions. Neural activity was recorded simultaneously from the antero-dorsal thalamic nucleus (AD) and post-subiculum (PoS) using chronically implanted, movable multi-site silicon probes. As the probes were advanced between sessions, we assumed that each session contained distinct units. From the total of 1,520 cells in these recordings, we used the criteria of the original study (concentration ratio, Raleigh test, and minimum firing rate threshold) to select 388 head direction cells.

Spike times (sampled at 20 kHz) were aligned to behavioural frames (39.0625 Hz) to determine head direction at each spike. The floating-point head direction at each timestamp was binned into 180 uniform bins spanning 0–360°. We created 6 repeats for each angle bin by dividing the timestamps into blocks of length ∼5 mins (12,000 frames), separated by 10 s intervals to avoid dependencies between consecutive blocks. Each block was randomly assigned to one of the six repeats, randomly discarding blocks if necessary to ensure each repeat contained the same number of blocks. A firing rate was then computed for each angle bin and repeat, as the number of spikes in all assigned frames divided by total length of these frames. The firing rate histograms from all cells of all recordings were put together to form a “virtual population” containing enough cells to form estimates of dimensionality.

### Regularized Cross-Validated PCA

We estimated the dimensionality of the visual cortical code to grating stimuli using an extension of the cross-validated Principal Components Analysis (cvPCA) method of Stringer et al. ^15^, which we term regularized cross-validated PCA (rcvPCA).

Ordinary PCA cannot be used for this purpose, as the variances it finds contain contributions from both stimulus-related (“signal”) variance, and stimulus-independent (“noise”) variance, but we are interested only in the former. The cvPCA method estimates the signal-related variance along a dimension **u** as a covariance between repeats 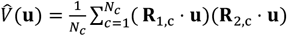 where **R**_i,c_ is the *N*_*θ*_ -dimensional vector of responses to repeat i of cell c across all stimuli. Because the stimulus-independent component of activity is independent across repeats, 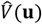 is an unbiased estimator of the stimulus-related variance along the direction **u** in stimulus space. Nevertheless, if the eigenvectors **u**_*n*_ are mis-estimated, even an unbiased estimate of the variance along these dimensions does not provide an accurate estimate of the stimulus-related principal component variances. In the original cvPCA method, stimuli were repeated twice, and the eigenvectors **u**_*n*_ were estimated by PCA of repeat 1 activity. The eigenvector estimates therefore contain contributions from both stimulus-dependent and stimulus-independent activity, and are not independent of the data used to assess variances, so may mis-estimate these eigenvectors.

The rcvPCA method improves over cvPCA by using three repeats of the stimuli. The first repeat is used to estimate the eigenvectors **u**_*n*_, using smoothing to obtain accurate estimates of stimulus-related covariance directions despite stimulus-independent variability. The second two repeats are used to estimate the stimulus-related variance along these eigenvectors, using the same formula 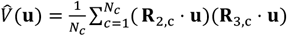 To avoid artifactually lowering the estimated dimensionality, smoothing is not used for repeats 2 and 3.

Although rcvPCA requires a minimum of 3 repeats, activity was initially binned and averaged to create 6 repeats per angular bin, as described above, for all datasets. This allowed multiple estimates of the eigenspectrum to be computed and averaged, thereby improving accuracy. We obtained our final eigenvalue estimates by averaging 60 runs of rcvPCA, one for each of the 6 ^5^*C*_2_ ways of assigning one of the 6 repeats to be the first rcvPCA repeat and two of the remaining 5 to be second and third.

#### Data Model

We model each response matrix as **R**_*i*_ = **Φ** + **E**_*i*_ for *i* = 1,2,3, where 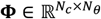 represents the reliable stimulus-related signal component, and **E**_*i*_ represents the noise component for repeat *i*. The noise matrices **E**_1_, **E**_2_, **E**_3_ are independent and identically distributed with mean zero.

#### Estimating eigenvectors from repeat 1

We estimate the eigenvectors by first computing smoothed tuning curves on repeat 1 and performing an SVD on the matrix of stacked tuning curves.

Let 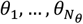 be the stimulus angles, and define the circular distance

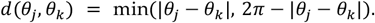

For *w* > 0, we define the smoothing kernel 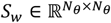 by a Gaussian:

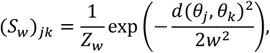

with *Z*_*w*_ chosen so that rows sum to one.

Define the smoothed training responses by 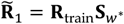 where *w*^∗^ is the optimal smoothing width determined by cross-validated least-squares. We define the stimulus eigenvector estimates 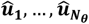 to be the L2-normalized eigenvectors of the covariance matrix 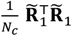

#### Unbiased Cross-Validated Eigenvalue Estimates

We estimate the stimulus-related variance in the *n*^*th*^ dimension as the covariance of population activity on repeats 2 and 3, projected onto the eigenvectors estimated from repeat 1:

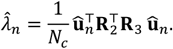

Since ***R***_2_ = **Φ** + ***E***_2_ and ***R***_3_ = **Φ** + ***E***_3_ with independent noise,

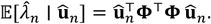

Thus 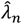 is an unbiased estimator of variance along û _*n*_, which will provide a good estimate of the true stimulus-related spectrum if the estimated eigenvector û_*n*_ is close to the true eigenvector. Note that the estimate 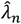 may be negative, even though the true PC variances cannot be, however this is required for unbiased estimation. Indeed, if the true variance were zero, an estimator that could only give positive values would necessarily be biased upward.

#### Validation

To validate that the rcvPCA method could indeed accurately estimate the PCA spectrum of stimulus-related neural activity, under conditions of trial-to-trial variability similar to those found in our recordings, we created a ground-truth dataset with identical variability to the real data, but for which the true spectrum was known (Fig. S1).

To do this we first estimated the noise of each cell, stimulus, and repeat, by subtracting from each cell’s response on each trial, its mean response over trials to the stimulus shown: 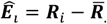. We then added this empirical noise to the deterministic stimulus response predicted by the AI model to create a simulation where the true stimulus-related activity was known. We found that rcvPCA found a spectrum that was very close to the spectrum computed by simple PCA on the noise-free stimulus responses (Fig. S1).

### Evaluating tuning curve equations by a single-cell level error

The AI equation discovery algorithm identified the best-performing orientation tuning function using only the Stringer et al. ^53^ data. To ensure that the model had validity for other datasets, we evaluated its predictive accuracy also on two other datasets collected in house with a 2-photon microscopy (conventional and light-beads). In all cases, single-cell error was computed using cross-validated explained variance (r^2^) relative to a Gaussian baseline.

Data were randomly partitioned along the stimulus axis into training, validation, and test sets. For each cell, the parameters of the AI and double-Gaussian models were fit on the training set by gradient descent, initialized at a start point generated by the AI-derived parameter estimator described below, stopping at the iteration when validation mean squared error no longer improved. Gradient descent was then re-run for the same number of iterations using the combined training and validation data. Performance was evaluated on the held-out test set through calculating the explained variance of each cell, 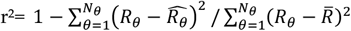 where 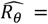 model prediction at angle i and 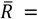 response mean across angles.

This procedure was applied identically to both the Gaussian and AI-derived models to ensure comparability. The resulting test r^2^ values provided a cross-validated measure of model fit.

### LLM-guided equation discovery

The equation discovery engine aims to find a simple formula that can accurately model each neuron’s tuning to sensory stimuli, using as few parameters as possible. It does this for a given experimental recording of neuronal responses to many stimulus presentations, represented as a matrix 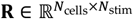, and a stimulus vector 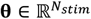 where *R*_*cs*_ is the response of cell *c* on trial *t* on which stimulus *θ*_*t*_ was presented.

The goal is to learn a generative neuron model *f*(*θ*; **β**) that predicts any cell’s mean response to any possible stimulus *θ*, given the cell’s tuning parameters 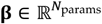 (e.g. preferred angle, baseline rate, amplitude, …). A candidate program *ϕ* produced by our system consists of 2 python functions:

1. The neuron model *f*(*θ*, **β**)

2. A parameter estimator 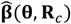 of the parameter vector 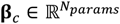 for each cell, given its responses 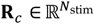 to stimuli 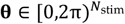. These estimated parameters are used as a starting point for gradient-based optimization.

The LLM–derived parameter estimator generates sensible initial values for the new tuning-curve model, thereby reducing the risk that gradient descent parameter estimation will converge to a local minimum, and thus allowing functions *f* whose loss landscapes are non-convex (Fig. S4). The resulting initial parameters substantially improved the performance and stability of subsequent gradient-based optimisation (Fig. 2e,f).

#### Core Operation: Stochastic Recombination Operator

The central operation in our evolutionary algorithm is a stochastic recombination operator ℛ that generates new programs from parent pairs. Given two parent programs *ϕ*_1_, *ϕ*_2_ sampled from the population and ranked by score such that *S*(*ϕ*_1_) ≤ *S*(*ϕ*_2_), the operator produces a new program:

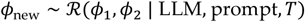

Where:

- LLM is a large language model (e.g., **Gemini 2.0 Flash**) that generates code
- prompt is template that a specifies how to create a prompt from parent programs, together with their scores and graphical diagnostics
- *T* is a temperature parameter which anneals over time, and controls the LLM’s stochasticity as well as the prompt text

The prompt is made up of the following components:

1. Temperature-dependent natural language instructions (if temperature is high, natural language instructions incentivize creativity, otherwise refinement is prioritized)
2. Source code of the two parent programs sorted by fitness (worst to best)
3. Diagnostic images showing parent model fits to data

The prompts follow the following format:

~~~
You are an AI scientist. Below are two programs sorted from worst to best. Your task is to generate a new mode l that is better than both of these.

[Temperature-dependent creativity instructions]

[Image analysis instructions]

Program 1 Score = S(ϕ1)
def neuron_model_v1(…): …
def param_estimator_v1(…): …

Program 2 Score = S(ϕ2)
def neuron_model_v2(…): …
def param_estimator_v2(…): …
~~~

Further details of the parts in square brackets are provided below. In practice, neuron models *f* and parameter estimators 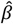 are generated in separate LLM calls, which we found empirically produces better output than asking an LLM to produce both functions simultaneously.

Examples of prompts, and code the LLMs generated in response to them, are in Supplementary Text 2.

#### Description of the winning model

The full python code of the winning model and its parameter estimator can be found in Supplementary Text 2. The winning model predicted each cell *c*’s response to stimulus *θ* as the sum of up to two asymmetric generalised Gaussian peaks:

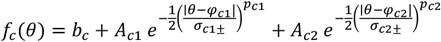

Here *b*_*c*_ is cell *c*’s baseline rate, *A*_*ci*_ its amplitude for peak *i, p*_*ci*_ the peakiness (shape) parameter, *ϕ*_*ci*_ the preferred angle, and *σ*_*ci*_ *±* the asymmetric widths. The symbols *σ*_*ci*_ *±* stand for two parameters *σ*_*ci*+_ and *σ*_*ci*−_ giving the widths to the left and right of the peak. In this notation,

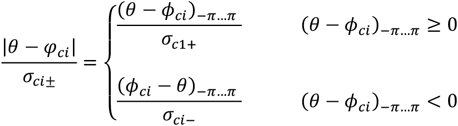

Where (*θ*)_−*π*…*π*_ means expressing the angle *θ* in the range (−*π, π*] by adding or subtracting multiples of 2*π* as necessary. The AI model thus has a total of 11 parameters per cell: one peakiness, amplitude, center, and two widths for each peak, plus one baseline rate.

#### Data Partitioning Strategy

Only cells with reliable responses to the grating stimuli were used to fit and evaluate tuning curves. For each cell c, we define the concentration and activity as follows:

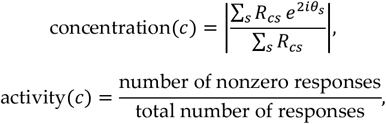

and only used cells with concentration ≥ 0.4 and activity ≥ 0.4.

To fit and evaluate the models, we employ a nested cross-validation scheme with two orthogonal splits:

#### Cell-Level Split (Outer Loop)

cells with reliable responses were divided into two groups:

- **Training cells** 𝒞_train_: 50% of cells (interleaved: *c*_1_, *c*_3_, *c*_5_, …) for program discovery.
- **Evaluation cells** 𝒞_test_: 50% of cells (interleaved: *c*_2_, *c*_4_, *c*_6_, …) reserved for final model selection and generalization assessment.

#### Trial-Level Split (Inner Loop)

- **Training trials** Θ_train_: 50% of stimulus presentations (interleaved: *θ*_1_, *θ*_3_, *θ*_5_, …) for parameter estimation and optimization
- **Testing trials** Θ_test_: 50% of stimulus presentations (interleaved: *θ*_2_, *θ*_4_, *θ*_6_, …) for scoring via cross-validated likelihood

#### Evaluation Protocol

- **During evolutionary search:** All programs *ϕ* are discovered using 𝒞_train_ only. For each training cell, parameters are estimated and optimized on Θ_train_, then evaluated on Θ_test_ to compute *S*(*ϕ*).
- **After search completion:** Top-scoring programs are evaluated on the held-out 𝒞_test_ using the same train-test split across trials. The program with best generalization to unseen cells is selected as the final model.

#### Program Scoring

To score a generated program 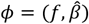, we first fit tuning parameters for each cell. Let 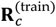 and 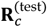 be the response vectors of cell *c* for the training and test trials. The program’s parameter estimator produces an initial parameter vector 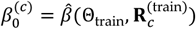, which is refined by gradient-descent (ADAM) to yield an estimate of:

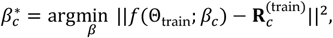

The program is then assigned a complexity-penalized, cross-validated score *S*(*ϕ*) by evaluating fit parameters across all cells on held-out test stimuli

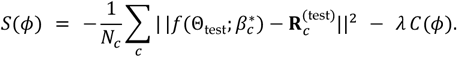

We set *C*(*ϕ*) to be the number of free parameters: *C*(*ϕ*) = dim(*β*_*c*_ ), and we include a small value of *λ* = 0.01 to prevent an explosion in the number of parameters without meaningful performance gains. Since the maximum observed reduction in held-out MSE was 0.55, each additional free parameter decreases the score by approximately 2% of the attainable improvement.

Because programs are LLM-generated, we enforce:

- **Type/shape checks** on outputs of the new *neuron_model* and *parameter_estimator*;

- The parameters introduced must be numeric, so they can optimized via gradient descent
- The parameter estimator must return an *n_train_cells x n_params* array
- The neuron model must return an *n_train_cells x n_train_stimuli* array
- **Timeouts** to discard programs that do not execute.

Programs that fail these checks receive a score of −∞ and are excluded from parent selection.

#### Horizontal Structure: Islands

We use a parallel island model with *N*_islands_ = 8 semi-independent populations evolving in parallel ^41^. Each island *j* maintains its own program pool 𝒫_*j*_ . New candidate programs are generated by selecting two parent programs *ϕ*_1_, *ϕ*_2_ uniformly from 𝒫_*j*_ and prompting the LLM to produce a child *ϕ*_new_ = ℛ(*ϕ*_1_, *ϕ*_2_). Each generation produces *B* = 6 new programs per island, for a total of 48 LLM calls per generation – chosen to stay within Gemini Tier 1 rate limits.

To balance exploration and exploitation, we implement migration after each generation. Each island *j* selects 2 of the newly generated programs to send to island (*j* + 1) mod *N*_islands_. Migrants are sampled according to a Boltzmann distribution with temperature *T* over program fitness:

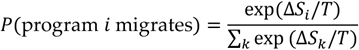

where Δ*S*_*i*_ is the normalized score difference from the island’s best program. At high temperatures (early iterations), migration is more uniform, allowing many programs to spread; at low temperatures, only elite programs migrate, accelerating convergence. Received migrants are integrated into the destination island’s population for recombination, and compete with local programs in the next pruning step.

#### Population Management and Deduplication

Each island *j* maintains a dynamic population 𝒫_*j*_ carrying out the following operations at the end of each generation:

1. **Deduplication**: Programs with high functional similarity (cosine similarity > 0.99 on evaluation matrices and score difference < 0.025) are removed to prevent genetic bottlenecks. Similarity is assessed both within islands and between island pairs—if two islands share ≥ 75% of the critical population size in duplicates, the island containing the lower scoring solutions is reinitialized with the seed models.
2. **Pruning** After each generation, populations are pruned by keeping only the 12 programs of highest score.

#### Temporal Structure: Evolution

Each island is initialized with **two seed programs**:

**Single-peaked Gaussian**:

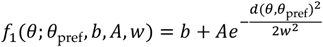

**Double-peaked Gaussian**:

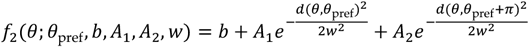

where

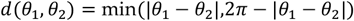

is the circular distance function.

Both programs include simple parameter estimators 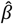 using circular statistics and peak detection on binned tuning curves, given in Supplementary Text 2.

The temperature is a global parameter across islands, which decays according to an annealing schedule of 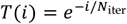, where *i* is the current generation. This temperature controls the evolution of programs via:

1. **LLM Temperature**: Controls the stochasticity of code generation
2. **Prompt Instructions**: Triggers a discrete phase transition from exploration mode to exploitation mode at a critical temperature *T*_*c*_
3. **Migration Dynamics**: Modulates the probability distribution governing inter-island program exchange

Full copies of the prompts can be found in Supplementary Text 2. Briefly, the *explore* mode instructs:

“Use the models below as inspiration, but be creative and invent something new. Which features correlate with lower loss? Find these features and extrapolate them. You should also combine features from several models, and experiment with new ideas.”

while the *exploit* mode emphasizes parsimony:

“Use the models below as a template. Which features correlate with lower loss? Focus on exploiting the strengths of existing models and eliminating their weaknesses or redundancies. Are the parameter ranges correct? You will be penalized for complexity, so make the new model as simple as possible while still being better.”

#### Programming language

To perform GPU-accelerated gradient descent we used JAX. However, we found during early development of the system that prompting the LLMs to generate code in JAX reduced creativity: LLM-generated JAX programs tended to use standard machine learning coding patterns whereas NumPy programs showed more scientifically creative solutions, perhaps due to the contents of the original LLM training data in these languages. We therefore prompted the LLMs to use NumPy syntax first, preserving creative LLM reasoning, then employed a second LLM to translate this code to JAX enabling to GPU-accelerated gradient-descent.

#### LLM Selection and Task Distribution

The system distributes computational load across three LLMs based on task complexity and creativity requirements:

- Lightweight Model (Default =*gemini-1*.*5-flash-8b*)

- Tasked with translating code from NumPy to JAX, following the main LLM call.
- Mid-Range Model (Default = *gemini-2*.*0-flash*)

- This reasonably lightweight model is much cheaper and faster than a thinking model, and we used it for 2/3 of the neuron models generated.
- We also used it to generate all parameter estimators, which only require heuristics. The prompts provided explicit constraints (≤100 lines, no complex fitting libraries like *scipy*.*optimize*).
- Thinking Model (Default = *gemini-2*.*5-flash* or *gemini-2*.*5-pro*)

- These were used to generate 1/3 of the neuron models, and were found to introduce more creative innovations. However, they were used sparingly as they are ∼3 times slower, and ∼20 times more expensive to run.

We reasoned that the mid-range model (*gemini-2*.*0-flash*) would produce modest but rapid refinements at volume, while periodic reasoning model injections would introduce occasional creative breakthroughs. This distribution of labor drastically reduces time and cost of programs generated. One run of the system that generated the program presented in the main text took 45 minutes of wall-clock time, and $8.25 of LLM tokens.

#### Multi-modal Prompt Engineering: Visual Feedback

In addition to the instructions and parent program code, the prompts contained a graphical input of diagnostic scatter plots, which showed each parent model’s predictions overlaid on raw neural data, for 9 randomly selected cells (Fig. S3). These images are embedded directly in the prompt alongside code, with instructions to:

- Identify systematic fit failures across multiple cells (e.g., *“model peaks are consistently too narrow in subplots 2, 5, 7”*)
- Analyze which model shapes (peak width, skewness, amplitude) best match data
- Propose parameter modifications addressing visual discrepancies Reference specific subplot examples in generated docstrings For example, prompts instruct:

*“Analyze the model’s fit to the data in the image below. Look for places where the model (red curve for best, green for second-best) deviates most from the binned data mean (blue curve). This is where the models are weakest, and where you should focus improvements. Do the models’ shapes align with the binned data mean and raw data scatter points? If not, how can you change the model to better match the data shape?”*

This visual feedback loop improves the convergence rate (Fig. 2f) and also provides insight into the reasoning the LLM used when crafting the models, which can be found in the docstrings of the LLM-generated code (Supplementary Text 2).

### Pseudo-Code for AI Equation Discovery System

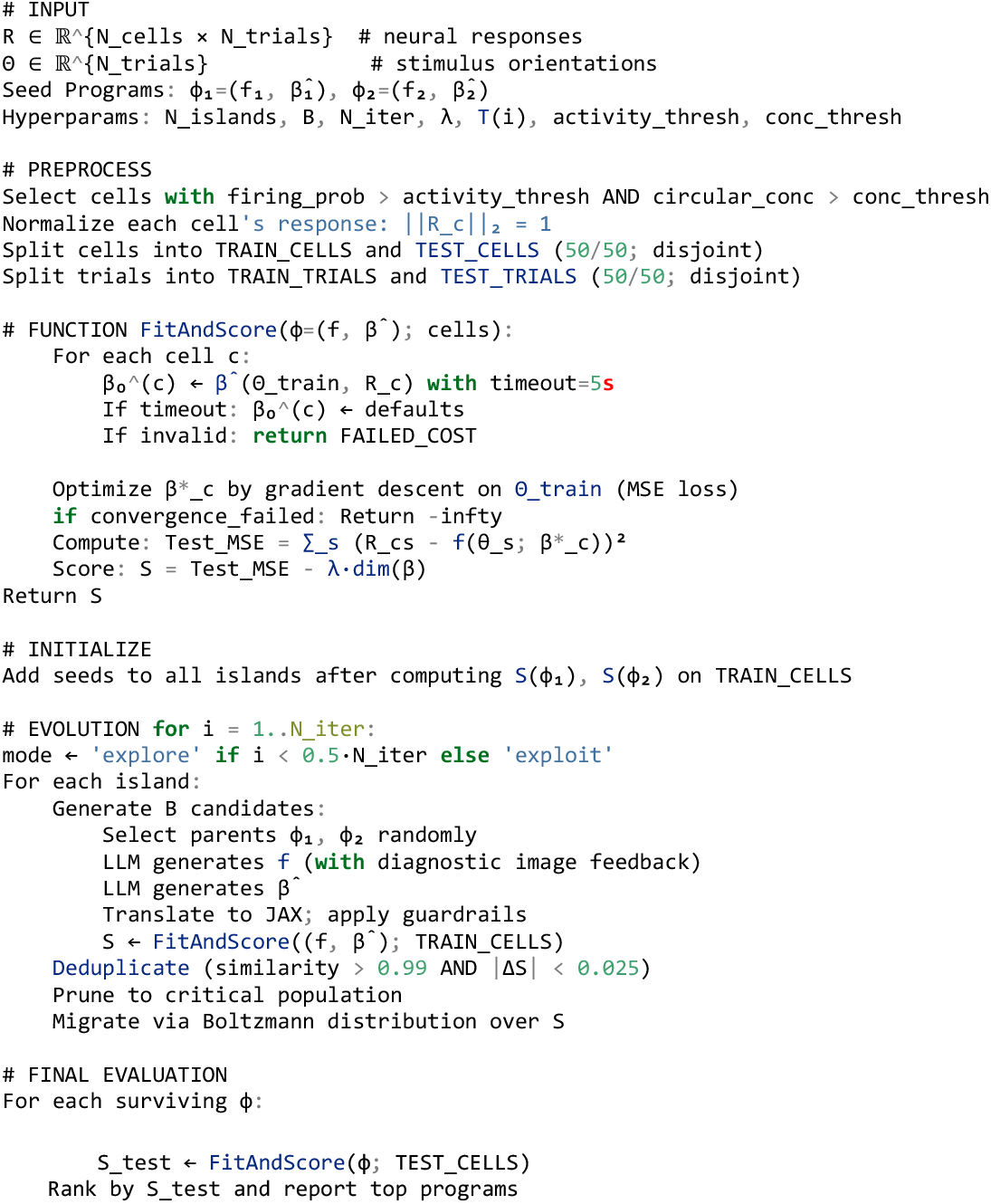

### Parameter ablation study

To assess how specific parameters of the AI model contributed to fit quality, we examined the effect on fit quality of successively ablating parameters (Fig. 3e). First, the full model (“AI model”) was fit and evaluated as described above. We next considered a model (“Gaussian+p+asym”) where the peak locations were constrained to be 180 degrees apart:

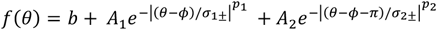

We fit the parameters of this model for each cell by gradient descent as described above, starting from the full-model fits of all parameters except ϕ, which was started at the location of the full model’s largest peak *ϕ*_1_.

The next ablated model (“Gaussian+p”) lost the asymmetric widths of the peaks, while allowing the two peaks to have different widths and peakiness:

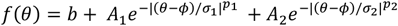

We again fit the parameters of this model by gradient descent, starting from the same parameters as (“Gaussian+p+asym”) except *σ*_1_ = (*σ*_1+_ + *σ*_1−_)/2 and *σ*_2_ = (*σ*_2+_ + *σ*_2−_)/2.

### Analysis of head direction cell tuning curves

Because head-direction cells, unlike orientation tuning curves, are single peaked, we modified the AI model to:

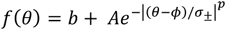

Visually observing the fit model also revealed that cells with highly asymmetric widths suffered from discontinuity at the angle antipodal to the cell’s preferred orientation. To correct for this discontinuity, we updated the tuning function by tiling the tuning functions either side of the original tuning function. Explicitly, for 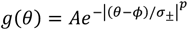, we define *g*_*ϕ*−2*π*_(*θ*) and *g*_*ϕ*+2*π*_, and let *f*(*θ*) = *g*_*ϕ*−2*π*_(*θ*) + *g*_*ϕ*_(*θ*) + *g*_*ϕ*+2*π*_(*θ*) for *θ* ∈ [0,2*π*]. Full and ablated models were fit as described above.

### Parameter estimation

The parameter-ablation and head-direction cell analyses of Figs. 3 and 4 required fitting the AI tuning curve model to neurons from all datasets, including head direction cells. To fit model parameters to these cells, we took the approach described above, avoiding local minima by using a gradient search starting point produced by the AI-generated parameter estimator. For this, we used parameter estimator code from a preliminary run of the AI system; the parameter estimator code found on the main run (shown in Supplementary Text 2.9) generates startpoints for both exponents as 2.0, which sufficed for it to discover the winning model, but is not ideal for scientific analysis. We now describe the algorithm this AI-generated code implements.

The parameter estimator first computes a smoothed firing rate histogram. To do so it divides the stimuli into *n = 180* uniform bins, computing summed activity and stimulus count histograms, convolving both with a Gaussian kernel (σ = 2.5), and dividing them to obtain a nonparametric tuning curve fit.

The baseline 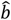 is estimated as the minimum of the smoothed curve. Local maxima are identified by comparing each bin to its immediate neighbours. Peaks are sorted by amplitude *Â*, with the largest designated as the primary peak. A secondary peak is included only if its amplitude exceeded 10% of the primary and its angular separation at least 90°. Cells without a second qualifying peak are treated as single-peaked (i.e. the second peak amplitude is set to 0).

To estimate the asymmetric widths *σ*_*i*_*±*, the parameter estimator identifies the half-height points between baseline and peak:

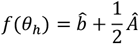

The corresponding angular distances to the peak, Δθ_*h*_, define the half-widths. Assuming a Gaussian form, the relationship between half-width and standard deviation is

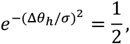

which yields

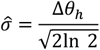

This procedure is applied independently to both sides of each peak to obtain asymmetric widths. Note that the LLM-generated code applies this formula even when *p* ≠ 2, but good results were still obtained.

To estimate the peakiness parameter *p*, the parameter estimator uses the ratio between the half-height (Δθ_*h*_ ) and quarter-height (Δθ_*q*_ ) angular distances. It estimates these using the formula

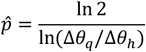

which can be obtained from the log of the generalized Gaussian −(Δ*θ*/*σ*)^*p*^ = ln *k*, taking the ratio for *k = ½* and *k = ¼*. This analytical relationship provides an efficient, interpretable estimate of p directly from the smoothed tuning curves.

### Readout stimulation

To investigate how high-dimensional coding arising from peaky tuning could enable linear readout, we simulated a hyperacuity task. A population of 3 60 neurons was simulated with preferred orientations *ϕ*_*i*_ spanning 360° at 1° intervals. The activity of each neuron was simulated with a single-peaked function:

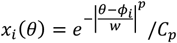

Where *w*=15°, *C*_*p*_ is a normalizing constant that ensures the mean-squared activity of all neurons is 1, *θ* is the current stimulus, and |*θ* − *ϕ*_*i*_| is shifted to like in the range -180… 180°. The target function was

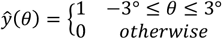

For delta-rule simulations (Fig. 5B), we used a logistic sigmoid output *y*(*θ*) = *σ*(***w*** ⋅ ***x***(*θ*) + *b*) and trained by gradient descent of the cross-entropy loss, i.e. the delta rule 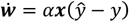, with a learning rate of *α*=0.1. Training was done in batches each containing 50 stimuli randomly uniformly sampled from -15°…15°. Performance was evaluated by cross-entropy loss of 300 evenly-sampled stimuli over the same range. Fig. 5B shows average performance over 20 runs of the simulation.

The delta rule is a local Hebbian rule as might be instantiated in a cortico-recipient brain structure such as the striatum, and its simulation was intended to capture how peaky tuning might benefit readout in such cases. However, it is not exactly solvable. To gain further insight we therefore considered a simpler, exactly-solvable case of mean-square error *E* = ∑_*θ*_ (*ŷ* (*θ*) − ***w*** ⋅ ***x***(*θ*))^2^. Finding the ***w*** that optimizes this under a constraint |***w***|^2^ = *A* is equivalent to solving the penalized minimization of *E* + *λ*|***w***|^2^. This is a ridge regression problem, which due to its convolution nature can be quickly solved by a frequency-domain Wiener filter:

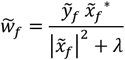

Here 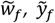 and 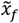 represent the Fourier transforms of the weight, target, and tuning curve vectors, and * represents complex conjugation.

The plot of Fig. 5d was obtained by running this Wiener filter for 4096 values of *λ* logarithmically spaced between 10^−8^ and 10^6^, and plotting the resulting weight norm vs. error; the value of *λ* resulting in weights closest to norm 1 was used to plot Fig. 5e,f.

### Mechanistic model simulation

The exactly solvable mechanistic model for generation of peaky tuning curves (illustrated in Fig. 6b-d) is described in Supplementary Text 1c. The present section contains a description of the code used to simulate this model, which produced the results in Fig. 6e-f. The simulations were run in custom python code.

101 excitatory pools were simulated, each by a single ordinary differential equation (ODE):

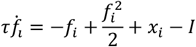

Where *τ* is the neuronal time constant of 10 ms; *x*_*i*_ is the external input received by pool *i*, simulated as a Gaussian function of *i* with center 0 and width 1, considering the 101 pools to be spread evenly over the range -2 to 2. The ODE was integrated for 300 ms with a 1 ms timestep, using scipy’s odeint function, and clipping activity to be non-negative. The inhibitory rate was an instantaneous function of excitatory activity, *I* = 75 ∑_*i*_(*x*_*i*_ − 1)_+_, where (*x*)_+_ denotes a ReLU function. There was no recurrent excitation.

**Figure S1.**
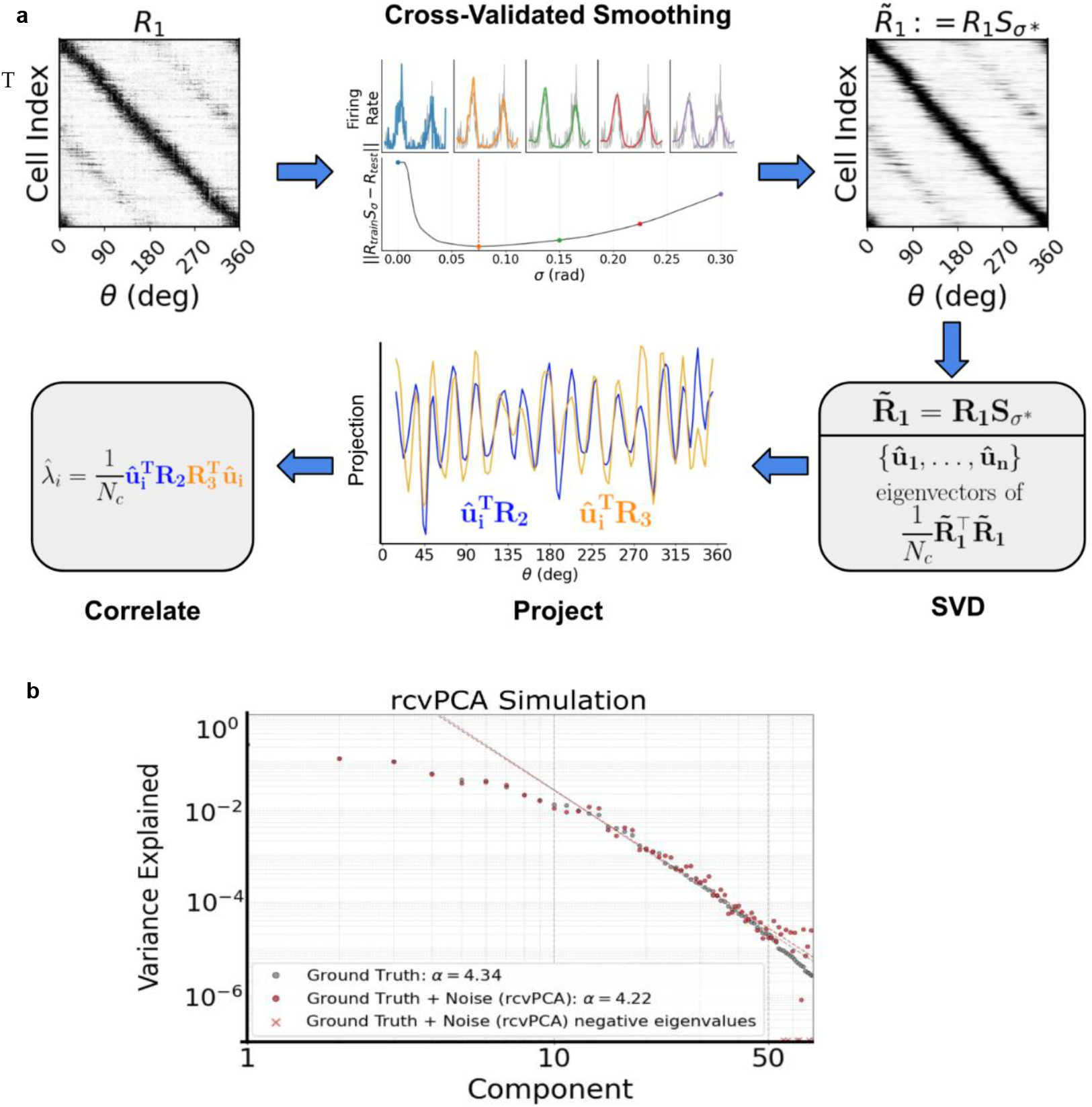
Regularized cross-validated principal component analysis (rcvPCA) **a**, Illustration of the rcvPCA method. The first repeat (R1) is used to nonparametrically estimate tuning curves for all cells by Gaussian smoothing. The smoothing width is determined by cross-validation to minimize prediction error, summed over cells, between the smoothed curves from R1 and responses on R2 and R3. Eigenvectors û_*i*_ are estimated by applying singular value decomposition (SVD) to the matrix of smoothed tuning curves. The remaining two repeats (R_2_ and R_3_) are projected onto these eigenvectors, and the cross-covariances of the projections provide estimates of the stimulus-related variance spectrum. Note that smoothing is not applied to R_2_ and R_3_ as this would artificially lower estimated dimensionality. **b**, Testing rcvPCA with ground-truth data. We simulated data in which the signal-related spectrum was known, and trial-to-trial variability matched that of the experimental data, by adding experimentally measured noise to a known ground-truth tuning spectrum. Noise for each cell *c*, stimulus θ, and repeat *i* was computed as the residual for repeat *i* relative to the repeat-averaged response: 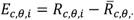, and added to a known tuning curve generated from the AI model’s fit to all cells: 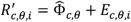. Gray points: PCA spectrum of the ground truth data, computed by standard PCA on the AI model tuning curves 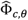. Gray dashed line: power-law fit to over eigenvalues 10-50, yielding an exponent of 4.34. Red points: spectrum estimated by rcvPCA from noisy data; red crosses are cases where the eigenvalue estimate was negative, which must be possible in order to have unbiased estimates of even strictly non-negative quantities. Red dashed line: power-law fit to rcvPCA, recovering a slope of 4.22, close to the ground truth value.

**Figure S2.**
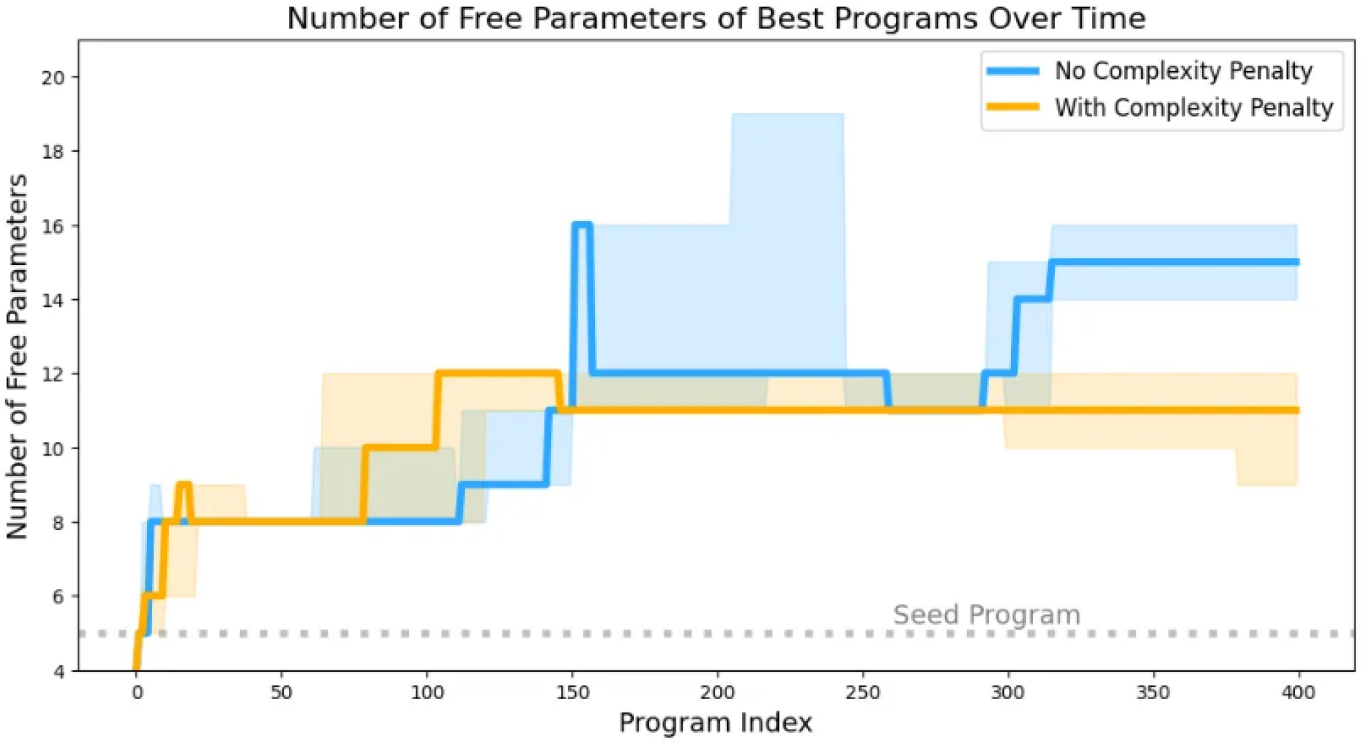
Effect of complexity penalty. The complexity penalty *λ* is proportional to the number of free parameters in the model. Setting λ = 0 (blue) causes the number of free parameters to grow over generations, yielding complex and opaque models. With the complexity penalty (orange), model size remains stable, models remain interpretable and overfitting to the training cells is controlled.

**Figure S3.**
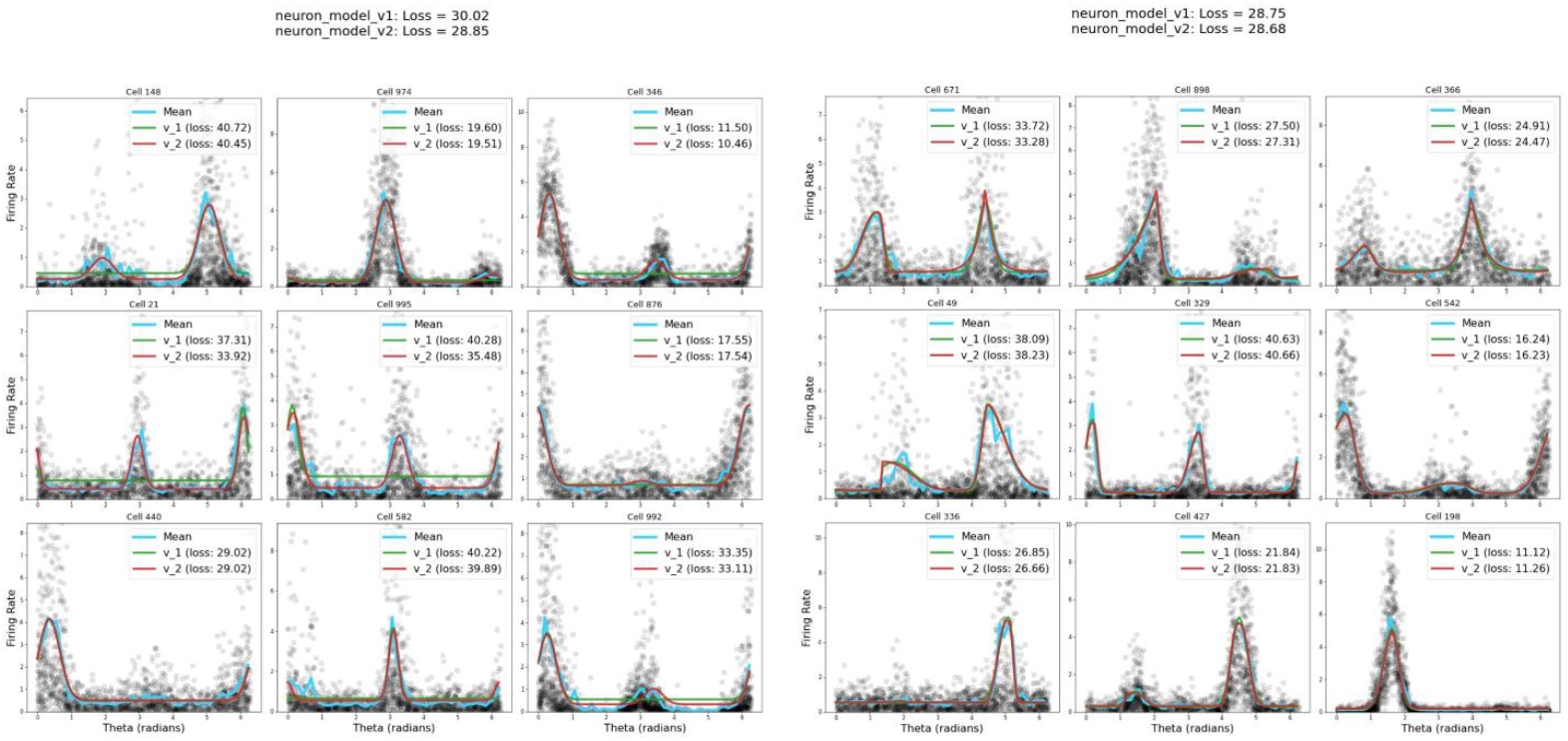
Graphical input to LLMs. In addition to textual prompts (described in Methods and quoted in full in Supplementary Text 2), the LLMs are given graphical diagnostics of parent model fits to 9 randomly-selected example cells. This figure shows the graphical input given to the LLM on the first iteration (left), and on the iteration that generated the winning model (right), at actual resolution. Black points show single-trial responses. Cyan curve shows binned-mean responses for 90 evenly-spaced bins. Red and green curves show fits of the two parent models. Note that the two examples show diagnostics for different randomly-selected cells.

**Figure S4.**
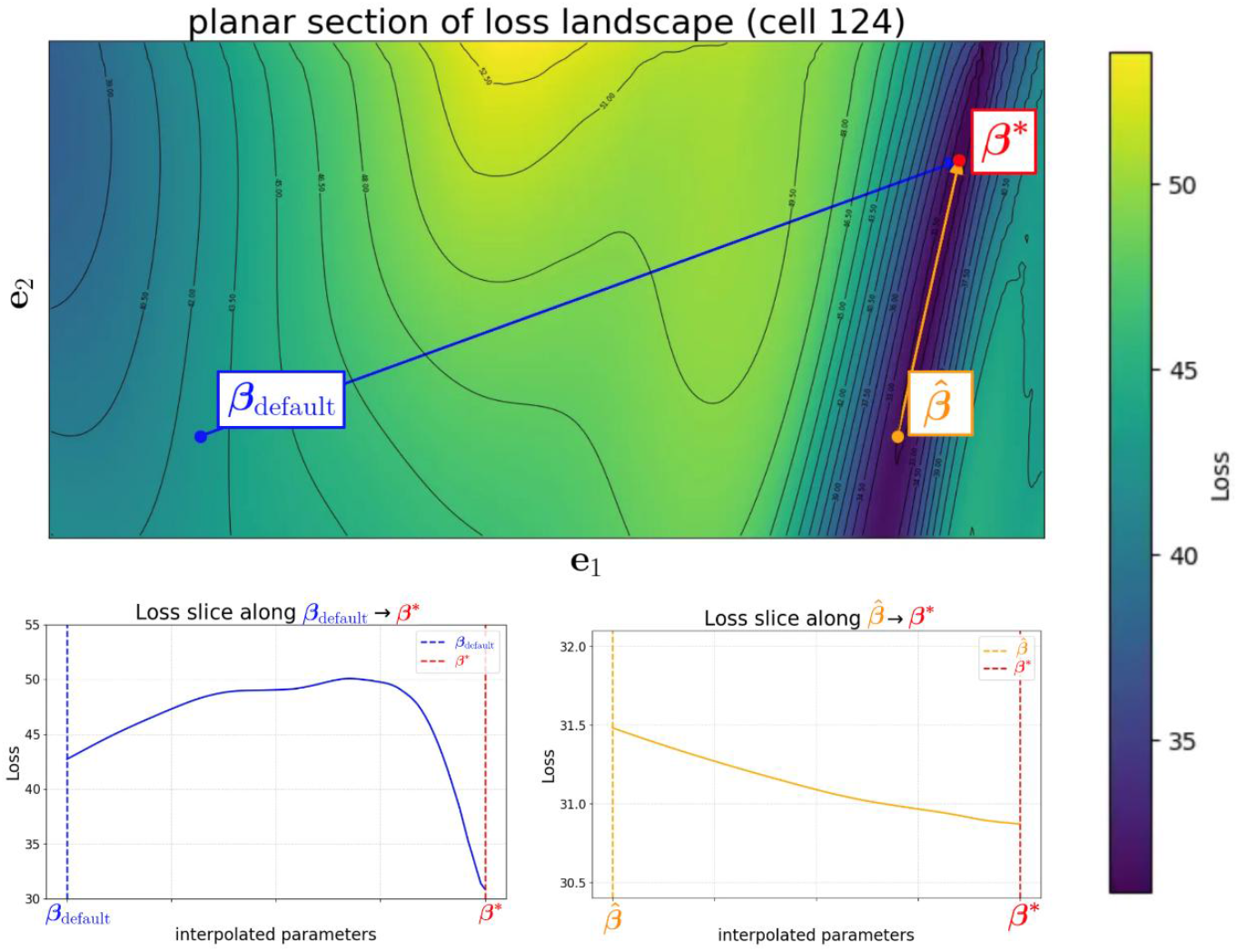
Avoiding local minima with the parameter estimator. Top: pseudocolor view of the winning AI model’s loss landscape for one cell. A planar section through the N-dimensional parameter space of the model is obtained by defining the x-axis by projection onto vector ***e***_**1**_ connecting the parameter estimator’s output 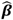 and a default parameter values ***β***_*de*f*ault*_ (defined by the LLM as the default parameters in the tuning function), and defining the y-axis by projection onto a vector ***e***_**2**_ orthogonally connecting the solution ***β***^**∗**^ to the line joining 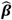 and ***β***_*de*f*ault*_. Bottom: projection of the loss landscape onto lines connecting the two possible start points. The line connecting ***β***_*de*f*ault*_ to ***β***^**∗**^ traverses a local maximum, but the line connecting 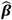 to ***β***^**∗**^ does not.

**Figure S5.**
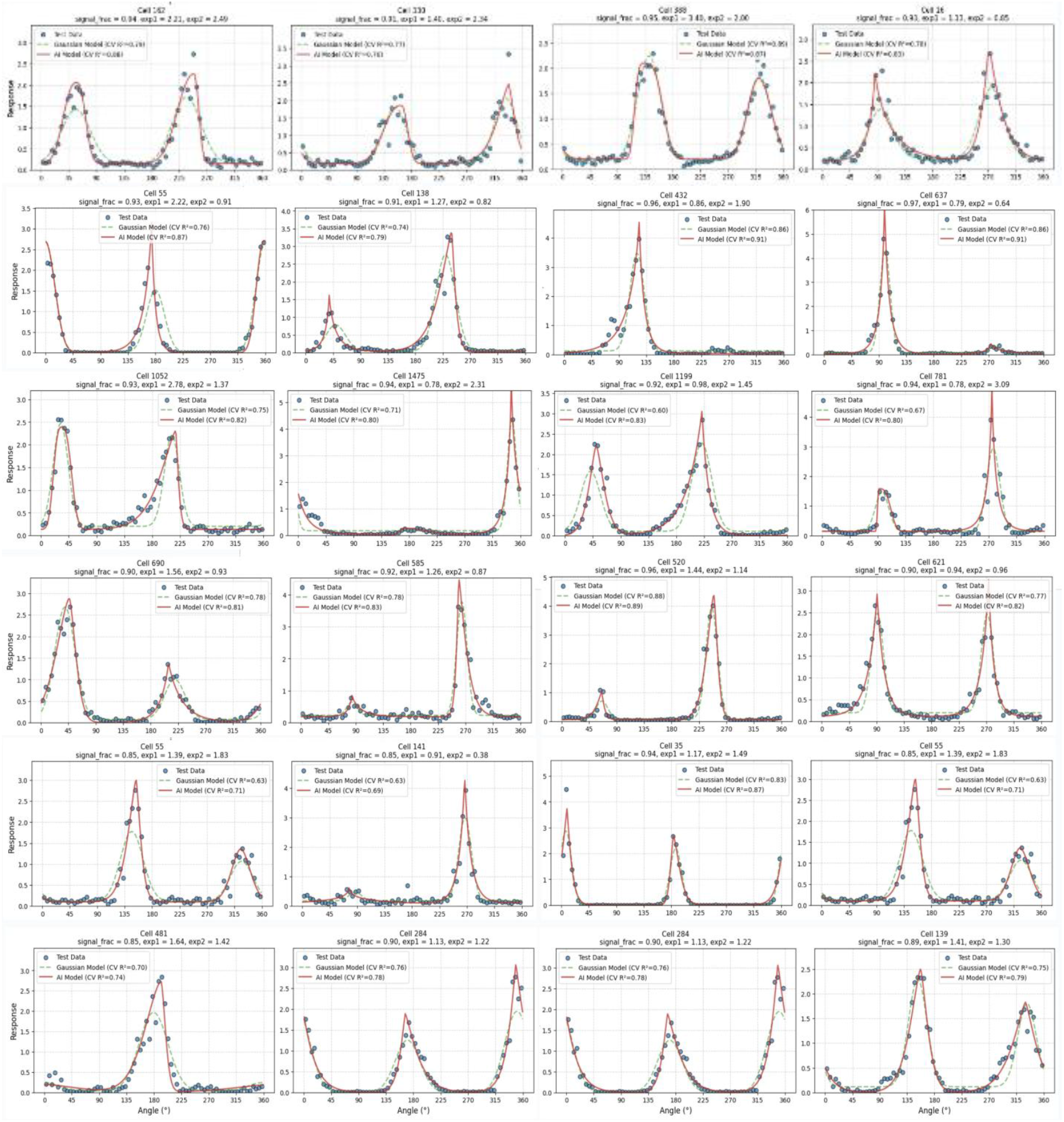
Tuning curve model fits for further example cells. Blue points: test-set responses as a function of grating direction for four example cells per mouse (one row per mouse; averaged over a random half the stimulus repeats, for any stimuli within the orientation bin). Red sold and green dashed curves show AI-derived and Gaussian model predictions, fit for each cell to training-set repeats.

**Figure S6.**
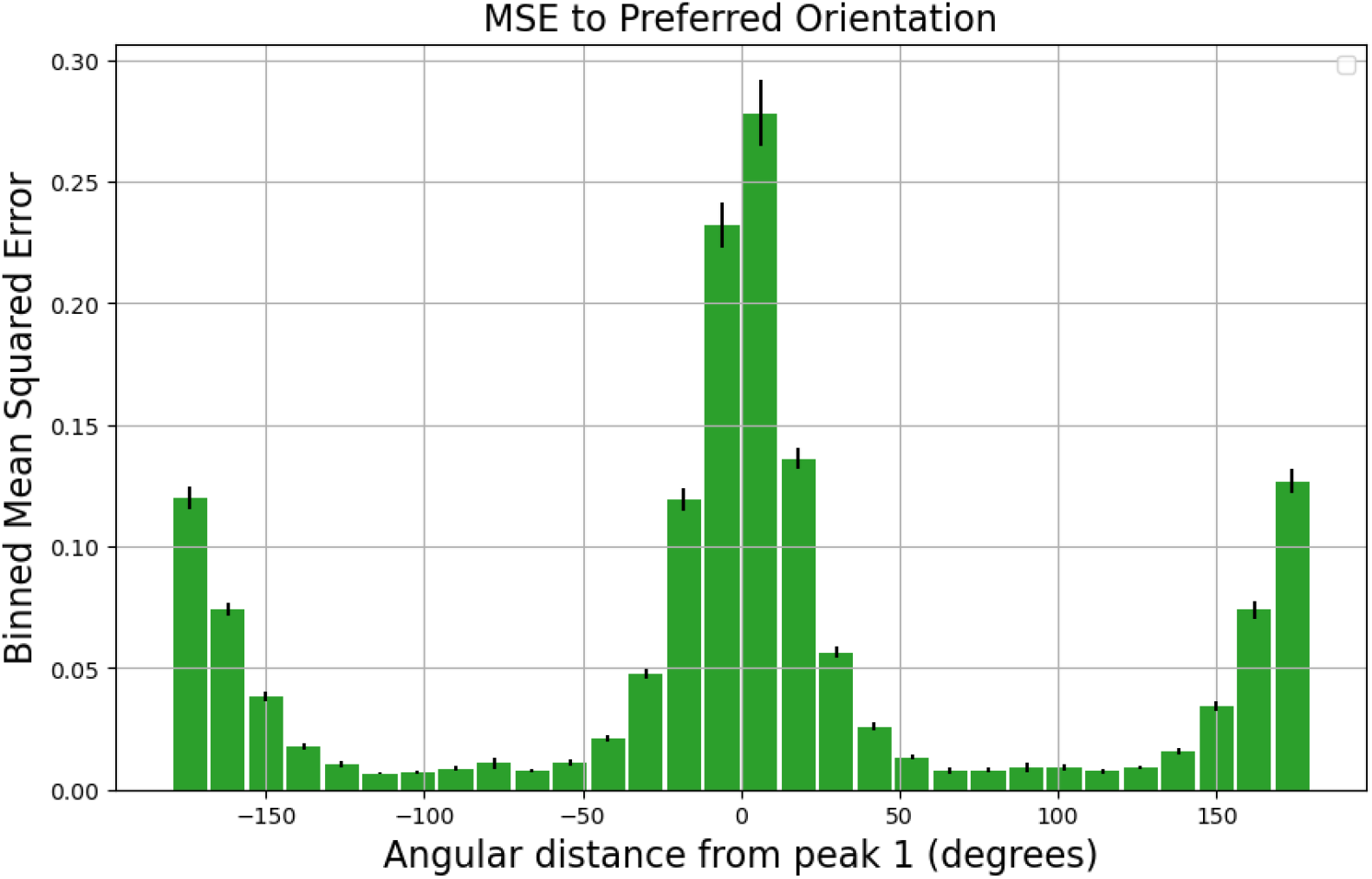
Gaussian model errors are at tuning curve peaks. Visual inspection of Gaussian model fits (Fig. 1b) suggested the Gaussian models failed to fit experimental data accurately specifically at the peaks of the tuning curves. To confirm this, we plotted the mean squared error of the Gaussian tuning curve fits as a function of angular distance from each cell’s preferred orientation for an example mouse. Large errors are concentrated near the tuning curve peaks, demonstrating that the Gaussian model failed specifically at its peaks. Error bars: standard error across cells.

**Figure S7.**
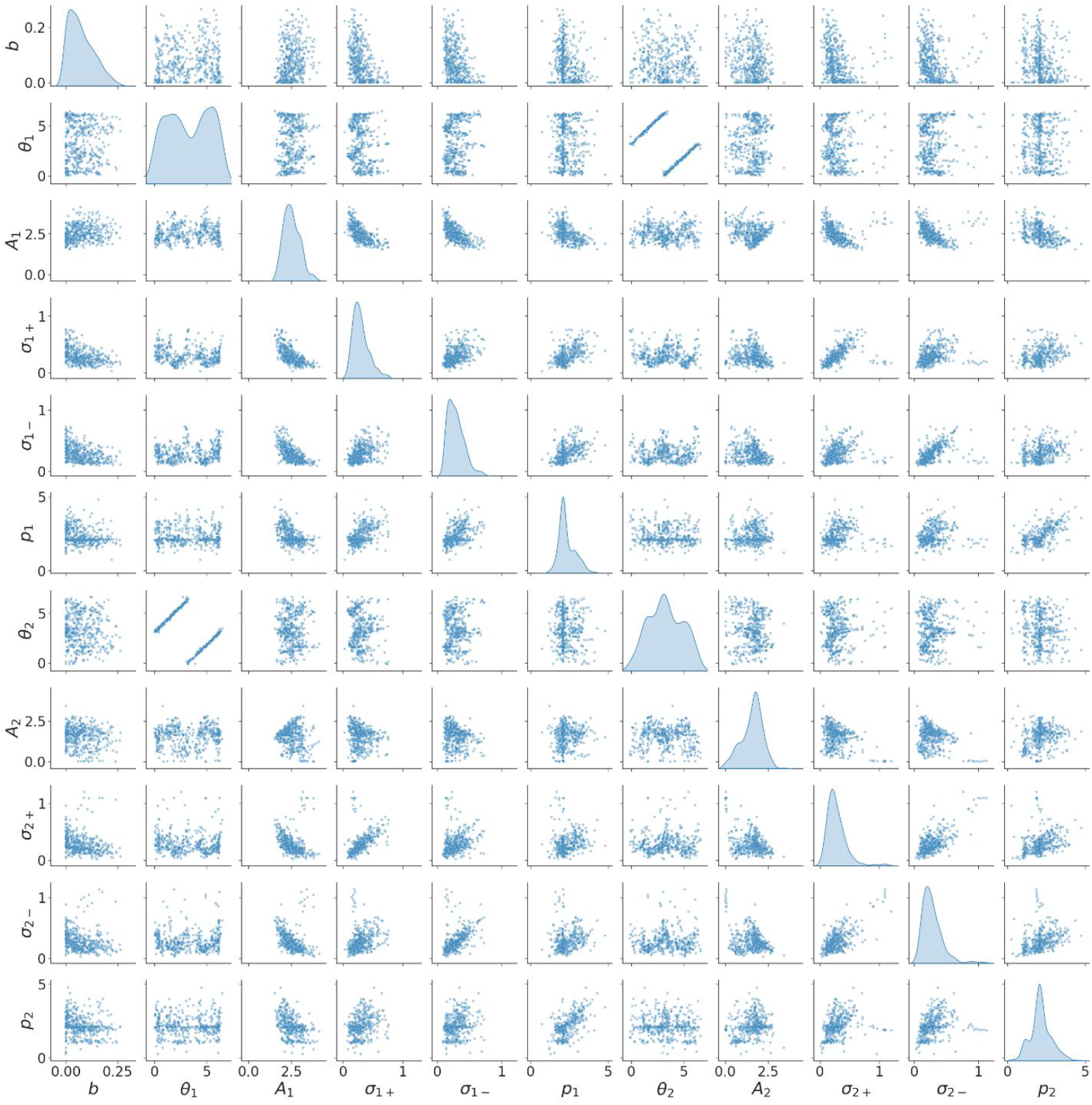
Scatterplot matrix showing distribution of parameters across cells for an example mouse. Each plot represents a pair of parameters, each point represents the parameters fit to one neuron. Five points corresponding to cells with outlying parameters are not visible on these axes.

## Supplementary Text 1a: Relationship of PCA and Fourier spectra

To understand the relationship between tuning curve shape and eigenspectrum, we consider a population of neurons with identically-shaped tuning curves *f*_*i*_(θ), with evenly-distributed preferred orientations *ϕ* ∈ [0,2*π*]. We let the size of the population tend to infinity, thus the activity of the population can be captured by a function of *ϕ*. For every θ, we represent this function of *ϕ* as a vector ***π*** in the Hilbert space *L*^2^([0,2*π*])

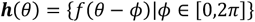

Whatever shape of the function *f*, the principal components of the population’s response to evenly distributed orientation stimuli are sine and cosine functions of *ϕ*. To show this we expand *f* in a Fourier cosine/sine series:

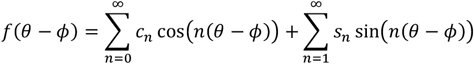

Note the cosine sum starts at *n* = 0 to include a constant. The projection of ***f*** onto a cosine basis cos(*mϕ*) is

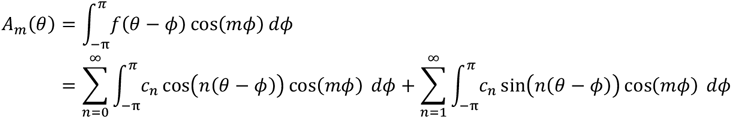

Using the trigonometric identities cos(*a*) cos(*b*) = (cos(*a* + *b*) + cos(*a* − *b*))/2 and cos(*a*) sin(*b*) = (cos(*a* + *b*) − cos(*a* − *b*))/2, we see that the only term that does not integrate to zero is

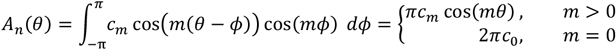

Thus the expected product of two projections across an ensemble of random stimuli is

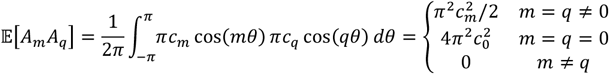

Similarly we obtain for the sine projections 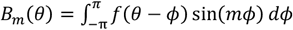

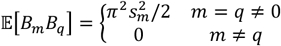

And the expectation of sine times cosine is zero

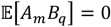

Thus, applying PCA to this population will yield eigenvectors whose weights on neurons with preferred orientation *ϕ* is given by sine or cosine functions of *ϕ*, with an eigenspectrum proportional to the corresponding squared Fourier components of the tuning function *f*(θ − *ϕ*).

## Supplementary Text 1b: Peak shape controls Fourier spectrum

To evaluate the eigenspectrum of our tuning functions, we must therefore calculate their Fourier series. We consider a simplified version of the tuning function with only one symmetrical peak:

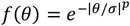

This function’s Fourier series is not exactly computable, but we can estimate the asymptotic decay of its Fourier spectrum using the relationship between a function’s smoothness and the asymptotic decay of its Fourier spectrum^1,2^. We will do so following the approach of Erdelyi^3^. Around θ= 0, we can expand in a power series:

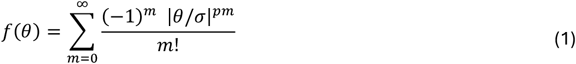

We will show the Fourier spectrum is dominated by the least-smooth term in this series, |θ/*σ*|^*p*^, which has Fourier decay *O*(*n*^−1−*p*^).

We can quantify a function’s smoothness by its differentiability: the more times it can be differentiated, the smoother it is. There are multiple measures of differentiability, for example whether we require that the derivative be continuous. Here we will use the concept of *weak differentiability*. A function *f*(*x*) has weak derivative *f*′(*x*) if we can integrate by parts: for any infinitely-differentiable *η*(*x*) with *η*(*a*) = *η*(*b*) = 0:

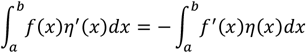

The weak derivative need not be continuous, or even defined at every point. For example, if *p* > 0 then |*x*|^*p*^ has weak derivative *p*|*x*|^*p*−1^sgn(*x*), which is not defined at 0 when *p* < 1, but can be still used for integration-by-parts. In general, |*x*|^*p*^ is weakly differentiable at least ⌊*p*⌋ + 1 times, where ⌊*p*⌋ is the largest integer ≤ *p*.

The Fourier series 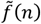 of a function *f*(θ) which is *m* times weakly differentiable must decay as *O*(*n*^−*m*^), i.e. there exists *C* > 0 such that 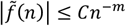 for all *n*. We show this by integrating by parts *m* times:

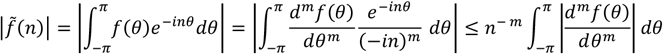

This means that the Fourier series of an infinitely differentiable function decays faster than any power; conversely, a function with power-law Fourier decay must have points where it is not infinitely differentiable.

Our tuning curve function 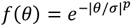 is infinitely differentiable except at two points: θ= 0, where 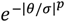 has a peak shaped like 1 − |θ/*σ*|^*p*^; and θ = *π*, where it has a weakly-differentiable “kink” due to the joining of the two decaying tails of the tuning curve wrapping on a circle. We can isolate the effect of these two points on the Fourier tail by considering the Fourier transform of *f*(θ)*η*(θ) + *f*(θ)*ζ*(θ), where *η*(θ) and *ζ*(θ) are suitable “bump functions”: infinitely-differentiable symmetrical functions such that *η*(θ) = 1, *ζ*(θ) = 0 for |θ| ≤ *A*, and *η*(θ) = 0, *ζ*(θ) = 1 for |θ| ≥ *B*, for some 0 ≤ *A* ≤ *B* ≤ *π*. Because *f*(θ) differs from *f*(θ)*η*(θ) + *f*(θ)*ζ*(θ) by an infinitely-differentiable function, the difference between their Fourier tails decays faster than any power.

We first consider *g*_*p*_(θ) = |θ/*σ*|^*p*^*η*(θ), which isolates the leading discontinuity at θ = 0. Because of the finite support and symmetry of *η*, we can simplify its Fourier integral:

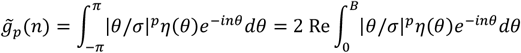

First note that for *p* > 0 we can integrate |θ/*σ*|^*p*^*η*(θ) against *e*^−*in*θ^ by parts ⌊*p*⌋ + 1 times to obtain 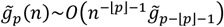. We thus need only prove that 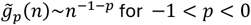.

We proceed by integrating |θ/*σ*|^*p*^ against *η*(θ)*e*^−*in*θ^ by parts. To do so we must first find an indefinite integral *G*(θ) with 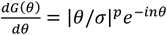. Then

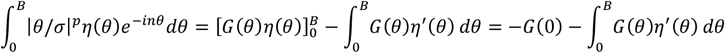

An indefinite integral suiting our needs is given by the complex contour integral:

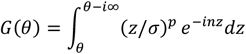

To show that 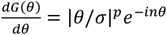, consider the contour connecting θ to the point at infinity. By Cauchy’s theorem, a contour deformation leaving the endpoints fixed and not crossing a singularity does not change the integral. Because there are no singularities away from *z* = 0, the contour can be deformed arbitrarily, so all dependence on θ comes from the lower integration limit. To compute 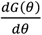, we can therefore simply differentiate with respect to the lower limit, which by the fundamental theorem of calculus gives 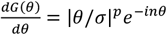. We will now use this indefinite integral *G*(θ) to bound our Fourier integral.

Making the substitution *z* = θ − *is*/*n* we get:

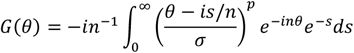

We can use this to obtain a bound on *G*:

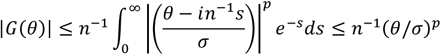

where we used the fact that |θ − *in*^−1^*s*| ≥ θ and *p* ≤ 0, so |θ − *in*^−1^*s*|^*p*^ ≤ θ^*p*^. Thus, we can bound the second term in the integration by parts,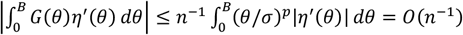.

We can also compute

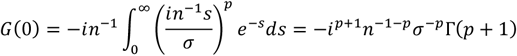

Thus

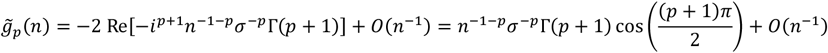

When *p* is not an even integer the cosine is non-zero, and 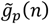 decays as *n*^−1−*p*^, which dominates over the *O*(*n*^−1^) term. This establishes that the Fourier series for *g*_*p*_(θ) = |θ/ *σ*|^*p*^*η*(θ) decays as *n*^−1−*p*^. Returning to the expansion (1), we see that the *m* = 1 term dominates, so the same is true for the original tuning curve 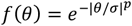.

It is also interesting to consider what happens as *p* approaches 2. So let *p* = 2 − *ϵ*. Then

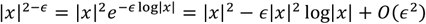

The first term is infinitely differentiable so its Fourier spectrum decays faster than any power, while the second term *ϵ*|*x*|^2^ log|*x*| has a Fourier spectrum *O*(*n*^−3^). Thus when *p* is close to 2, its Fourier tail does decay as a power law with exponent *n*^−1−*p*^, but the amplitude of this tail goes to zero as *p* → 2. This can also be seen from the cos 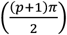 term in the formula for 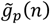, which approaches 0 as p gets close to an even integer.

The AI tuning equation contains two terms 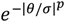, and the Fourier series will be dominated by the peak with smaller *p*. Thus, the eigenspectrum of the AI model decays as *n*^−*α*^, where *α* = −2(1 + min(*p*_1_, *p*_2_)).

## Supplementary Text 1c: Mechanistic model of tuning curve peaks

We consider a network with pools of recurrently connected excitatory neurons, each maximally tuned to one possible stimulus (in simulations we used 101 possible stimuli arranged linearly, with the actual stimulus in the center), and global feedback inhibition (Fig. 6a). The excitation within a pool is supralinear, so the network is a simple variant of the “stabilized supralinear network” of Rubin et al.^4^, with local excitation and global inhibition. Network dynamics results in a “soft winner-take-all” effect, that transforms smooth tuning curves to sharp peaks.

First consider one excitatory pool, receiving external excitatory input *x*_i_ and global feedback inhibition *I*. We approximate all neurons in the pool as having the same rate *f*_*i*_. We model the dynamics of this pool as:

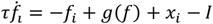

Here *g*(*f*) is a supralinear function, which without loss of generality we take to be 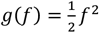; because the transformation of the tuning curve peak shape depends only on local properties of *g*(*f*), this covers any case with a 2^nd^ order Taylor expansion. The steady-state solution satisfies:

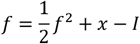

For 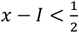 this has two solutions,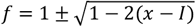, of which the lower is stable and the upper unstable. For 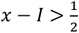, there is no stable solution, and *f* increases without bound. Thus, for stable activation, we must have 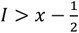.

Now consider a network comprising N such local excitatory pools with external inputs *x*_i_ and rates *f*_*i*_, and one global inhibitory pool with rate *I*. The activity of the inhibitory pool is determined by the total excitatory input it receives: *I* = ∑_i_ *h*(*f*_*i*_). We consider the function *h* to have a high gain and a threshold at 1. With such negative feedback, the system will converge to a state where none of the *f*_*i*_ are substantially above 1, the threshold at which inhibition becomes very strong, but at least one of them is close to it. For simulations we will use a ReLU for *h*, but the model is exactly solvable for a step function (*h* = 0 for *f* < 1 and *h* = ∞ for *f* ≥ 1) in which the stable state exactly satisfies 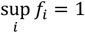 . Substituting into the equation 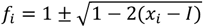, we get 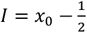, where 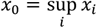 . Inhibition constrains every pool’s activity to be *f*_*i*_ ≤ 1, and thus below the unstable fixed point, so every pool converges to the steady-state fixed point:

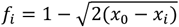

Finally, let us consider how this transforms tuning curves. Suppose that the input *x*_i_ depends smoothly on a parameter, for example Gaussian tuning depending on the proximity of a sensory stimulus θ to the pool’s preferred stimulus *ϕ*_*i*_. Because we are interested in the transformation of the peak we may approximate the sensory tuning with a 2^nd^ order Taylor series: *x*_i_ = *A*(1 − (*ϕ*_*i*_ − θ)^2^ + *O*(*ϕ*_*i*_ − θ)^4^). The neuron receiving maximal input will be the with preferred stimulus θ, thus *x*_0_ = *A*. So, we have

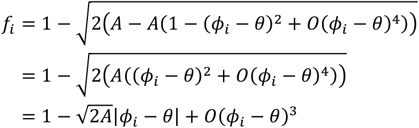

This analysis therefore predicts that the network transforms an infinitely-differentiable Gaussian-tuned synaptic input into a firing rate output with a sharp peak proportional to the absolute value function |θ − *ϕ*_*i*_|.

A simulation of this network, using a ReLU for the function *h* instead of a step function, confirmed that the steady-state activity of the network indeed converted a Gaussian-tuned synaptic input to a sharply-peaked firing rate output (Figure 6e-f). Another notable feature of the network’s dynamics was that it showed progressive sharpening of tuning curves, a phenomenon which has been observed in the visual and auditory cortices ^5–7^.

## Supplementary Text 2: Example prompts and discovered programs

This appendix quotes examples of the prompts given to the LLMs, and the outputs produced in response. It shows:

1. The prompt used to evolve new neuron models on the first generation. This includes a graphical diagnostic, text explaining the task required of the LLM, and code for the two (human-written) seed models. In early iterations, the prompt has “explore”-mode instructions.
2. The new neuron model produced by the LLM in response to this prompt. Note that the docstring explains the LLM’s reasoning for producing this model, referencing the peak positions in the graphical diagnostic.
3. The prompt used to create the parameter estimator for the above model. This contains text explaining the task required of the LLM, code for the two seed models and their (human-written) parameter estimators, and code for the new neuron model whose parameter estimator is required
4. The parameter estimator code produced in response to this prompt.
5. The prompt used to translate the neuron model produced on the first iteration from numpy to JAX. The parallelization offered by JAX is required for gradient ascent, but we found that asking LLMs to write code in JAX resulted in less creative solutions.
6. The JAX code produced in response to this prompt
7. The prompt that generated the winning neuron model. This has a similar form to the first iteration, but contains “exploit”-mode instructions.
8. The neuron model produced in response to this prompt, which instantiates the winning model.
9. The parameter estimator code produced for the winning model.

### 1. Prompt for neuron model (Generation 1)

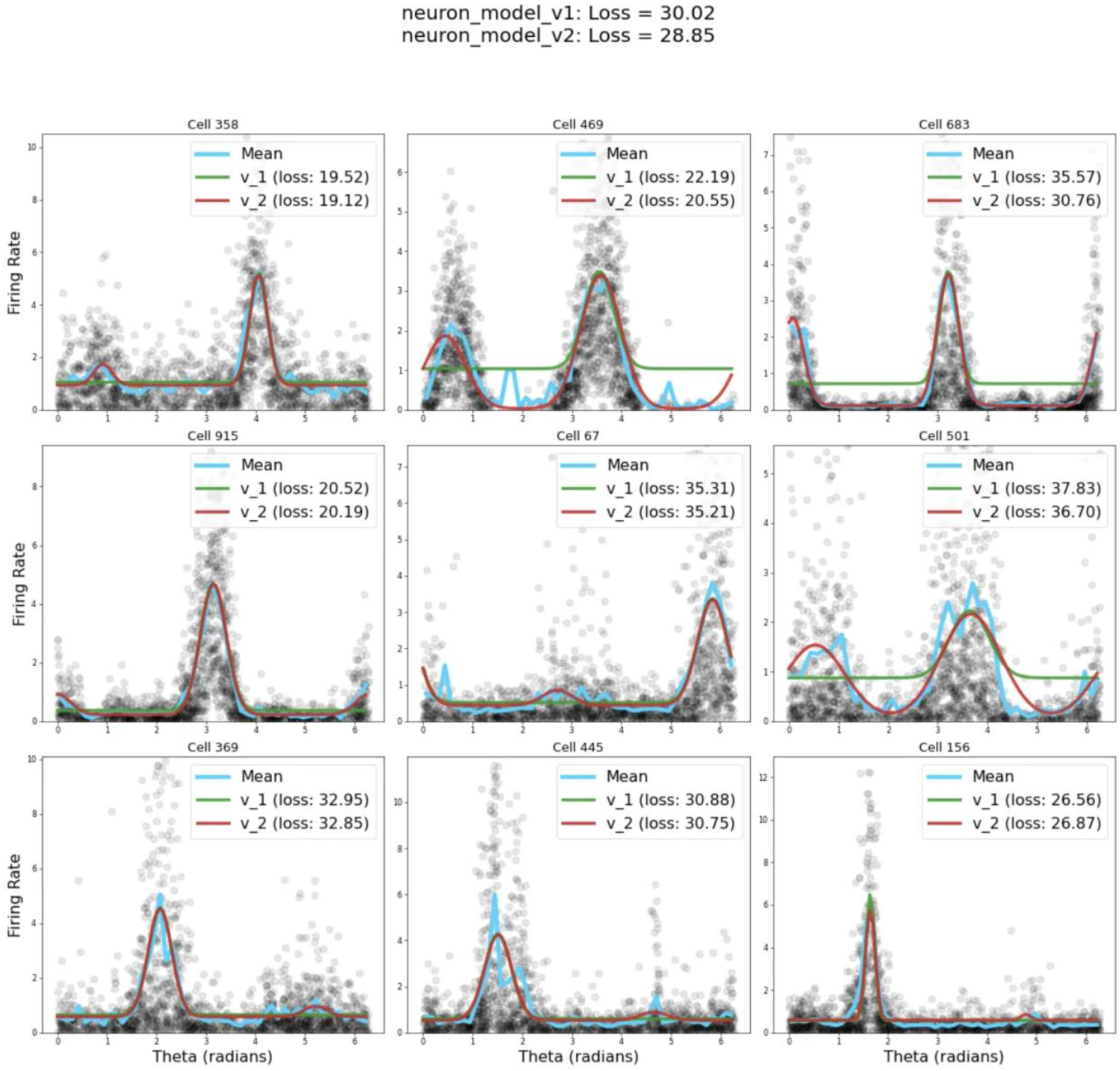

~~~
You are an AI scientist. The programs below are biological models of neurons. The models are sorted from highest to lowest loss.

Your task is to create a new neuron model, neuron_model_v3, that has a lower loss than the models below.

*Analyze* the progression of the models, *generalize* the improvements, and *create* a new model that is better than *all* previous models.

Use the models below as inspiration, but be *creative* and *invent* something new. Which features in the models below correlate with lower loss? Find these features and
*extrapolate* them. You should also *combine* features from several models, and
*experiment* with new ideas.

**Image Analysis Instructions:**

Attached is a scatter plot of the neuron models’ performance on top of raw neural data. The binned mean is plotted in **sky-blue**, ‵neuron_model_v1‵ is plotted in **green**, and ‵neuron_model_v2‵ is plotted in **red**.
Analyse the models’ fits to the data in the image below. Identify systematic weaknesses of the models by observing patterns across multiple cell plots. For instance, consider: 
*  **Model Comparisons:** Which models are better for each cell? That is to say, which models track the blue curve better? Which features of the models are responsible for improving the fit?
*  *    Model Fit:** How well do the models fit the binned data mean? Look for places where even the models (**red** curve for best model, **green** for second best model) deviate most from the binned data mean (**blue** curve). This is where the models are weakest, and where you should focus your improvements.
**    Model Shape:** Do the models’ shapes (e.g., peak sharpness, width, skewness, amplitude, etc.) align with the binned data mean (**blue**) and raw data scatter points (**black**)? If not, how do they differ? How can you change the model to better match the data shape?
*  **Parameter Flexibility:** Are there free parameters that could be introduced or modified to better capture the observed response profiles? Utilize your analysis of the shortcomings of the current models’ shapes and add free parameters or modify existing ones to address these issues.

Use this analysis to inform the design of a new neuron model, ‵neuron_model_v3‵, that improves upon the previous models.

Include your analysis of the image in the docstring of your new model. Point to specific subplots in the image that illustrate the *strengths* and *weaknesses* of the parent models. Explain how you plan to **fix** the weaknesses of the parent models.

**Code Generation Guidelines:**

* Import any packages you use.
* Do not include any text other than the code.
* Ensure all free parameters are numeric, not strings.
* At the beginning of the code, clip the free parameters to a biologically plausible range, e.g., ‵theta_pref = np.clip(theta_pref, 0, 2 * jnp.pi)‵.

**Docstring Guidelines:**
* Begin by listing the parent models and give them a name that describes their key features, e.g., ‵parent_model_1: simple_exponential_decay-model‵, ‵parent_model_2: double_exponential_decay_model‵. Never refer to the models as ‵neuron_model_v1‵, ‵neuron_model_v2‵, etc. Instead, refer to them as ‵parent_models‵ or their descriptive names (e.g. ‵simple_exponential_decay_model‵).
* Do not refer to the current model as ‵neuron_model_v3‵. Instead, refer to it as “this model”.
* Provide a simple equation for the model, including all free parameters.
* Include a brief description of how the model improves upon the previous models, citing specific features or changes that lead to lower loss.
~~~

**`loss of model 1: 30.02`**

~~~
import numpy as np
def neuron_model_v1(theta, theta_pref=0.0, baseline=0.0, amplitude=1.0, tuning_width=1.0):
    “““
    A simple neuron model that computes the response based on a Gaussian tuning curve.
    Args:
        theta (np.ndarray): The angle in radians.
        theta_pref (float): Preferred direction of the neuron.
        baseline (float): Baseline firing rate.
        amplitude (float): Maximum firing rate above baseline.
        tuning_width (float): Width of the tuning curve.
    Returns:
        np.ndarray: The firing rate of the neuron at angle theta.
    “““
        theta_pref = np.clip(theta_pref, 0, 2 * np.pi)
        baseline = np.clip(baseline, 0, None)
        amplitude = np.clip(amplitude, 0, None)
        tuning_width = np.clip(tuning_width, 0.01, None)
        circ_dist_rad = lambda theta1, theta2: np.abs(np.arctan2(np.sin(theta1 - theta2), np.cos(theta1 - theta2)))
        dist = circ_dist_rad(theta, theta_pref)
        return baseline + amplitude * np.exp(-0.5 * (dist / tuning_width) ** 2)
~~~

**`loss of model 2: 28.85`**

~~~
import numpy as np
def neuron_model_v2(theta, theta_pref=0.0, baseline=0.0, amplitude_1=1.0, amplitude_2=0.0,
tuning_width=1.0):
    “““
    A neuron model that computes the response based on a double peaked gaussian tuning curve, with peaks at theta_pref and (theta_pref + pi) % 2pi.
    Args:
        theta (np.ndarray): Input angles in radians.
        theta_pref (float): Preferred angle in radians.
        baseline (float): Baseline firing rate.
        amplitude_1 (float): Amplitude of the first peak.
        amplitude_2_ratio (float): Ratio of the second peak’s amplitude to the first peak’s
amplitude.
        tuning_width (float): Width of the tuning curves around preferred angles.
    Returns:
        np.ndarray: The response of the neuron model.
    “““
        theta_pref = np.clip(theta_pref, 0, 2 * np.pi)
        baseline = np.clip(baseline, 0, None)
        amplitude_1 = np.clip(amplitude_1, 0, None)
        amplitude_2 = np.clip(amplitude_2, 0, None)
        tuning_width = np.clip(tuning_width, 0.01, None)
        circ_dist_rad = lambda theta1, theta2: np.abs(np.arctan2(np.sin(theta1 - theta2), np.cos(theta1 - theta2)))
        dist_1 = circ_dist_rad(theta, theta_pref)
        dist_2 = circ_dist_rad(theta, (theta_pref + np.pi) % (2 * np.pi))
        return baseline + amplitude_1 * np.exp(-0.5 * (dist_1 / tuning_width) ** 2) + amplitude_2 * np.exp(-0.5 * (dist_2 / tuning_width) ** 2)
~~~

### 2. Output for neuron model (Generation 1)

~~~
import numpy as np
def neuron_model(theta, theta_pref=0.0, baseline=0.0, amplitude_1=1.0,
tuning_width_1_left=1.0, tuning_width_1_right=1.0, amplitude_2=0.0,
tuning_width_2_left=1.0, tuning_width_2_right=1.0):
    “““
    parent_model_1: simple_gaussian_tuning_model
    parent_model_2: double_gaussian_tuning_model

    Equation for this model:
    R(theta) = baseline +
               amplitude_1 * exp(-0.5 * (abs(signed_dist_1) / sigma_1_eff)^2) +
               amplitude_2 * exp(-0.5 * (abs(signed_dist_2) / sigma_2_eff)^2)
    where:
    signed_dist_1 = arctan2(sin(theta - theta_pref), cos(theta - theta_pref))
    sigma_1_eff = tuning_width_1_left if signed_dist_1 < 0 else tuning_width_1_right
    signed_dist_2 = arctan2(sin(theta - (theta_pref + pi)), cos(theta - (theta_pref + pi)))
    sigma_2_eff = tuning_width_2_left if signed_dist_2 < 0 else tuning_width_2_right

    This model improves upon previous models by providing greater flexibility in tuning curve shapes, particularly by
    allowing for asymmetric Gaussian peaks and independent widths for the primary and secondary peaks.
    The ‘simple_gaussian_tuning_model’ (neuron_model_v1) often failed to capture neurons with bi-directional tuning,
    resulting in high loss for cells like Cell 469, Cell 683, Cell 67, and Cell 501, where it only fits one peak and
    a flat baseline for the rest of the circle (e.g., green line for Cell 469 missing the peak around 4.5 rad).

    The ‘double_gaussian_tuning_model’ (neuron_model_v2) was a significant improvement, introducing a second peak at
    the anti-preferred direction. This dramatically reduced loss by fitting the second peak observed in the data mean
    (blue line) for many cells (e.g., comparing red vs green lines for Cell 358, Cell 469, Cell 683, Cell 915).
    Despite this, the ‘double_gaussian_tuning_model’ assumed a shared and symmetric tuning_width for both peaks, which
    is often not true in biological data.

    This model addresses the limitations of the ‘double_gaussian_tuning_model’ by:
    1. Allowing the preferred (amplitude_1) and anti-preferred (amplitude_2) peaks to have *independent* widths.
       Even if ‵tuning_width_1_left‵ and ‵tuning_width_1_right‵ are equal (for symmetry), and similarly for
       the second peak, this model permits ‵tuning_width_1_left‵ (or
‵tuning_width_1_right‵) to be different from
       ‵tuning_width_2_left‵ (or ‵tuning_width_2_right‵). This enhanced flexibility can be beneficial where the
       main peak is broader or sharper than the anti-preferred peak, as can be visually suggested in Cell 67
       or Cell 469 where the peaks visually appear to have different extents.
    2. Introducing *asymmetry* to each Gaussian peak, using separate width parameters for the left and right
       sides of each peak (e.g., tuning_width_1_left and tuning_width_1_right for the first peak). This is a crucial
       feature for fitting real neural tuning curves, which are rarely perfectly symmetric
Gaussians. For example,
       in Cell 469, the main peak (around 1.5 rad) of the ‘Mean’ (blue curve) appears to
have a steeper rise and a
       shallower decay, a characteristic not well-captured by the symmetric red ‘v2’ curve. Similarly, in Cell 501,
       the peak around 4 rad might benefit from asymmetric fitting to better match the blue mean curve.
       By modeling this asymmetry, this model can more precisely track the observed firing rate profiles, leading to
       a lower overall loss by minimizing systematic deviations between the model and the data’s true shape.
    Args:
       theta (np.ndarray): The angle in radians for which to compute the neuron’s response.
       theta_pref (float): The preferred direction of the neuron for the primary peak.
       baseline (float): The baseline (minimum) firing rate of the neuron.
       amplitude_1 (float): The maximum firing rate above baseline for the primary peak. tuning_width_1_left (float): The tuning width for the ‘left’ side (counter-clockwise) of the primary peak.
       tuning_width_1_right (float): The tuning width for the ‘right’ side (clockwise) of the primary peak.
       amplitude_2 (float): The maximum firing rate above baseline for the secondary peak (at theta_pref + pi).
       tuning_width_2_left (float): The tuning width for the ‘left’ side (counter-clockwise) of the secondary peak.
       tuning_width_2_right (float): The tuning width for the ‘right’ side (clockwise) of the secondary peak.
    Returns:
       np.ndarray: The firing rate of the neuron at angle theta.
    “““
    # Clip parameters to biologically plausible ranges
    theta_pref = np.clip(theta_pref, 0, 2 * np.pi)
    baseline = np.clip(baseline, 0, None) # Firing rate cannot be negative
    amplitude_1 = np.clip(amplitude_1, 0, None)
    amplitude_2 = np.clip(amplitude_2, 0, None)

    # Tuning widths must be positive and non-zero to avoid division by zero
    tuning_width_1_left = np.clip(tuning_width_1_left, 0.01, None)
    tuning_width_1_right = np.clip(tuning_width_1_right, 0.01, None)
    tuning_width_2_left = np.clip(tuning_width_2_left, 0.01, None)
    tuning_width_2_right = np.clip(tuning_width_2_right, 0.01, None)

    # Calculate signed circular distance for the first peak
    signed_dist_1 = np.arctan2(np.sin(theta - theta_pref), np.cos(theta - theta_pref))

    # Determine effective tuning width for the first peak based on signed distance
    sigma_1_eff = np.where(signed_dist_1 < 0, tuning_width_1_left, tuning_width_1_right)

    # Calculate response for the first peak
    peak_1_response = amplitude_1 * np.exp(-0.5 * (np.abs(signed_dist_1) / sigma_1_eff) **
2)
    # Calculate signed circular distance for the second peak (anti-preferred direction)
    theta_antipref = (theta_pref + np.pi) % (2 * np.pi)
    signed_dist_2 = np.arctan2(np.sin(theta - theta_antipref), np.cos(theta - theta_antipref))

    # Determine effective tuning width for the second peak based on signed distance
    sigma_2_eff = np.where(signed_dist_2 < 0, tuning_width_2_left, tuning_width_2_right)

    # Calculate response for the second peak
    peak_2_response = amplitude_2 * np.exp(-0.5 * (np.abs(signed_dist_2) / sigma_2_eff) **
2)
    return baseline + peak_1_response + peak_2_response
~~~

### 3. Prompt for parameter estimator (Generation 1)

~~~
You are an AI scientist. Your task is to create a simple parameter estimator function, parameter_estimator_v3, to estimate the free parameters of the latest neuron model, neuron_model_v3.

The parameter should be estimated directly, using statistical principles and knowledge of what the parameters represent biologically.
*Analyze* the progression of the parameter estimators, *generalize* the improvements, and
*create* a new parameter estimator that is better than *all* previous estimators.
**Code Generation Guidelines:**
* Import any packages you use.
* Do not include any text other than the code.
* The only arguments to the function should be the stimuli and the spike count.
* Your response **must** be less than {max_lines} lines (including imports). If it is longer, it will be immediately rejected.
* Do not attempt to fit the parameters using complex fitting functions like ‵curve_fit‵, ‵least_squares‵ or ‵minimize‵. This function should be a simple starting point for the parameter estimation.
~~~

**`loss of model 1: 30.02`**

~~~
import numpy as np
def neuron_model_v1(theta, theta_pref=0.0, baseline=0.0, amplitude=1.0, tuning_width=1.0):
    “““
    A simple neuron model that computes the response based on a Gaussian tuning curve. 
    Args:
        theta (np.ndarray): The angle in radians.
        theta_pref (float): Preferred direction of the neuron.
        baseline (float): Baseline firing rate.
        amplitude (float): Maximum firing rate above baseline.
        tuning_width (float): Width of the tuning curve.
    Returns:
        np.ndarray: The firing rate of the neuron at angle theta.
    “““
    theta_pref = np.clip(theta_pref, 0, 2 * np.pi)
    baseline = np.clip(baseline, 0, None)
    amplitude = np.clip(amplitude, 0, None)
    tuning_width = np.clip(tuning_width, 0.01, None)
    circ_dist_rad = lambda theta1, theta2: np.abs(np.arctan2(np.sin(theta1 - theta2),
                                                                 np.cos(theta1 - theta2)))
    dist = circ_dist_rad(theta, theta_pref)
    return baseline + amplitude * np.exp(-0.5 * (dist / tuning_width) ** 2)
def parameter_estimator_v1(theta, spike_counts):
    “““
    Estimates the parameters of the gaussian neuron model. We do this by creating a binned
       tuning curve and picking out salient features.
    Args:
       theta (np.ndarray): Angles in radians.
       spike_counts (np.ndarray): Spike counts corresponding to each angle.
    Returns:
       np.ndarray: Estimated parameters [theta_pref, baseline, amplitude, tuning_width].
    “““
    n_bins = 20
    bin_idx = ((theta * n_bins) / (2 * np.pi)).astype(np.int32)
    bin_idx = np.clip(bin_idx, 0, n_bins - 1)
    sums = np.bincount(bin_idx, weights=spike_counts, minlength=n_bins)
    counts = np.bincount(bin_idx, minlength=n_bins)
    tuning_curve = np.zeros(n_bins, dtype=np.float32)
    tuning_curve[counts > 0] = sums[counts > 0] / counts[counts > 0]
    pref_idx = np.argmax(tuning_curve)
    theta_pref = pref_idx * (2 * np.pi / n_bins)
    baseline = np.min(tuning_curve)
    amplitude = np.max(tuning_curve) – baseline
    half_max = baseline + amplitude / 2.0
    indices = (np.arange(-5, 6) + pref_idx) % n_bins
    above_half_max = tuning_curve[indices] >= half_max
    full_width_half_max = 2 * np.pi * np.sum(above_half_max) / n_bins
    tuning_width = full_width_half_max / (2.0 * np.sqrt(2 * np.log(2)))
    return np.array([theta_pref, baseline, amplitude, tuning_width])
~~~

**`loss of model 2: 28.85`**

~~~
import numpy as np
def neuron_model_v2(theta, theta_pref=0.0, baseline=0.0, amplitude_1=1.0, amplitude_2=0.0,
tuning_width=1.0):
   “““
   A neuron model that computes the response based on a double peaked gaussian tuning
       curve, with peaks at theta_pref and (theta_pref + pi) % 2pi.
   Args:
       theta (np.ndarray): Input angles in radians.
       theta_pref (float): Preferred angle in radians.
       baseline (float): Baseline firing rate.
       amplitude_1 (float): Amplitude of the first peak.
       amplitude_2_ratio (float): Ratio of the second peak’s amplitude to the first peak’s amplitude.
       tuning_width (float): Width of the tuning curves around preferred angles.
   Returns:
       np.ndarray: The response of the neuron model.
   “““
       theta_pref = np.clip(theta_pref, 0, 2 * np.pi)
       baseline = np.clip(baseline, 0, None)
       amplitude_1 = np.clip(amplitude_1, 0, None)
       amplitude_2 = np.clip(amplitude_2, 0, None)
       tuning_width = np.clip(tuning_width, 0.01, None)
       circ_dist_rad = lambda theta1, theta2: np.abs(np.arctan2(np.sin(theta1 - theta2),
                                                                    np.cos(theta1 - theta2)))
       dist_1 = circ_dist_rad(theta, theta_pref)
       dist_2 = circ_dist_rad(theta, (theta_pref + np.pi) % (2 * np.pi))
       return baseline + amplitude_1 * np.exp(-0.5 * (dist_1 / tuning_width) ** 2) +
       amplitude_2 * np.exp(-0.5 * (dist_2 / tuning_width) ** 2)
def parameter_estimator_v2(theta, spike_counts):
   “““
   A parameter estimator for the double peaked neuron model. Creates a binned tuning curve from spike counts and estimates parameters using features from the tuning curve.
   Args:
       theta (np.ndarray): Input angles in radians. (n_trials,)
       spike_counts (np.ndarray): Spike counts corresponding to the angles. (n_trials,)
   Returns:
       np.ndarray: Estimated parameters [theta_pref, baseline, amplitude_1, amplitude_2_ratio, tuning_width].
   “““
   n_bins = 50
   bin_idx = ((theta * n_bins) / (2 * np.pi)).astype(np.int32)
   bin_idx = np.clip(bin_idx, 0, n_bins - 1)
   sums = np.bincount(bin_idx, weights=spike_counts, minlength=n_bins)
   counts = np.bincount(bin_idx, minlength=n_bins)
   def gaussian_kernel(sig: int) -> np.ndarray:
       x = np.arange(-int(3 * sig), int(3 * sig) + 1)
       k = np.exp(-0.5 * (x / sig) ** 2)
       return k / np.sum(k)
   k = gaussian_kernel(2)
   pad = len(k) // 2
   sums_padded = np.pad(sums, (pad, pad), mode=‘wrap’)
   counts_padded = np.pad(counts, (pad, pad), mode=‘wrap’)
   num_conv = np.convolve(sums_padded, k, mode=‘valid’)
   den_conv = np.convolve(counts_padded, k, mode=‘valid’)
   tuning_curve = num_conv / (den_conv + 1e-8)
   pref_idx = np.argmax(tuning_curve)
   theta_pref = pref_idx * (2 * np.pi / n_bins)
   baseline = np.min(tuning_curve)
   amplitude_1 = np.max(tuning_curve) - baseline
   amplitude_2 = tuning_curve[(pref_idx + n_bins // 2) % n_bins] – baseline
   half_max = baseline + amplitude_1 / 2.0
   indices = (np.arange(-5, 6) + pref_idx) % n_bins
   above_half_max = tuning_curve[indices] >= half_max
   full_width_half_max = 2 * np.pi * np.sum(above_half_max) / n_bins
   tuning_width = full_width_half_max / (2.0 * np.sqrt(2 * np.log(2)))
   return np.array([theta_pref, baseline, amplitude_1, amplitude_2, tuning_width])
import numpy as np
def neuron_model_v3(theta, theta_pref=0.0, baseline=0.0,
amplitude_1=1.0, tuning_width_1_left=1.0,
tuning_width_1_right=1.0, amplitude_2=0.0, tuning_width_2_left=1.0, tuning_width_2_right=1.0):
   “““
   parent_model_1: simple_gaussian_tuning_model parent_model_2: double_gaussian_tuning_model
   Equation for this model:
   R(theta) = baseline +
              amplitude_1 * exp(-0.5 * (abs(signed_dist_1) / sigma_1_eff)^2) +
              amplitude_2 * exp(-0.5 * (abs(signed_dist_2) / sigma_2_eff)^2)
   where:
   signed_dist_1 = arctan2(sin(theta - theta_pref), cos(theta - theta_pref))
   sigma_1_eff = tuning_width_1_left if signed_dist_1 < 0 else tuning_width_1_right
   signed_dist_2 = arctan2(sin(theta - (theta_pref + pi)), cos(theta - (theta_pref + pi)))
   sigma_2_eff = tuning_width_2_left if signed_dist_2 < 0 else tuning_width_2_right
   This model improves upon previous models by providing greater flexibility in tuning curve shapes, particularly by
   allowing for asymmetric Gaussian peaks and independent widths for the primary and secondary peaks.
   The ‘simple_gaussian_tuning_model’ (neuron_model_v1) often failed to capture neurons with bi-directional tuning,
   resulting in high loss for cells like Cell 469, Cell 683, Cell 67, and Cell 501, where it only fits one peak and
   a flat baseline for the rest of the circle (e.g., green line for Cell 469 missing the peak around 4.5 rad).
   The ‘double_gaussian_tuning_model’ (neuron_model_v2) was a significant improvement, introducing a second peak at
   the anti-preferred direction. This dramatically reduced loss by fitting the second peak observed in the data mean
   (blue line) for many cells (e.g., comparing red vs green lines for Cell 358, Cell 469, Cell 683, Cell 915).
   Despite this, the ‘double_gaussian_tuning_model’ assumed a shared and symmetric tuning_width for both peaks, which
   is often not true in biological data.
   This model addresses the limitations of the ‘double_gaussian_tuning_model’ by:
   1. Allowing the preferred (amplitude_1) and anti-preferred (amplitude_2) peaks to have *independent* widths.
      Even if ‵tuning_width_1_left‵ and ‵tuning_width_1_right‵ are equal (for symmetry), and similarly for
      the second peak, this model permits ‵tuning_width_1_left‵ (or ‵tuning_width_1_right‵) to be different from
      ‵tuning_width_2_left‵ (or ‵tuning_width_2_right‵). This enhanced flexibility can be beneficial where the
      main peak is broader or sharper than the anti-preferred peak, as can be visually suggested in Cell 67
      or Cell 469 where the peaks visually appear to have different extents.
   2. Introducing *asymmetry* to each Gaussian peak, using separate width parameters for the left and right
      sides of each peak (e.g., tuning_width_1_left and tuning_width_1_right for the first peak). This is a crucial
      feature for fitting real neural tuning curves, which are rarely perfectly symmetric Gaussians. For example,
      in Cell 469, the main peak (around 1.5 rad) of the ‘Mean’ (blue curve) appears to have a steeper rise and a
      shallower decay, a characteristic not well-captured by the symmetric red ‘v2’ curve. Similarly, in Cell 501,
      the peak around 4 rad might benefit from asymmetric fitting to better match the blue mean curve.
      By modeling this asymmetry, this model can more precisely track the observed firing rate profiles, leading to
      a lower overall loss by minimizing systematic deviations between the model and the data’s true shape.
   Args:
      theta (np.ndarray): The angle in radians for which to compute the neuron’s response.
      theta_pref (float): The preferred direction of the neuron for the primary peak.
      baseline (float): The baseline (minimum) firing rate of the neuron.
      amplitude_1 (float): The maximum firing rate above baseline for the primary peak.
      tuning_width_1_left (float): The tuning width for the ‘left’ side (counter-
clockwise) of the primary peak.
      tuning_width_1_right (float): The tuning width for the ‘right’ side (clockwise) of the primary peak.
      amplitude_2 (float): The maximum firing rate above baseline for the secondary peak (at theta_pref + pi).
      tuning_width_2_left (float): The tuning width for the ‘left’ side (counter-clockwise) of the secondary peak.
      tuning_width_2_right (float): The tuning width for the ‘right’ side (clockwise) of the secondary peak.
   Returns:
      np.ndarray: The firing rate of the neuron at angle theta.
   “““
   # Clip parameters to biologically plausible ranges
   theta_pref = np.clip(theta_pref, 0, 2 * np.pi)
   baseline = np.clip(baseline, 0, None) # Firing rate cannot be negative
   amplitude_1 = np.clip(amplitude_1, 0, None)
   amplitude_2 = np.clip(amplitude_2, 0, None)
   # Tuning widths must be positive and non-zero to avoid division by zero
   tuning_width_1_left = np.clip(tuning_width_1_left, 0.01, None)
   tuning_width_1_right = np.clip(tuning_width_1_right, 0.01, None)
   tuning_width_2_left = np.clip(tuning_width_2_left, 0.01, None)
   tuning_width_2_right = np.clip(tuning_width_2_right, 0.01, None)
   # Calculate signed circular distance for the first peak
   signed_dist_1 = np.arctan2(np.sin(theta - theta_pref), np.cos(theta - theta_pref))
   # Determine effective tuning width for the first peak based on signed distance
   sigma_1_eff = np.where(signed_dist_1 < 0, tuning_width_1_left, tuning_width_1_right)
   # Calculate response for the first peak
   peak_1_response = amplitude_1 * np.exp(-0.5 * (np.abs(signed_dist_1) / sigma_1_eff) ** 
2)
   # Calculate signed circular distance for the second peak (anti-preferred direction)
   theta_antipref = (theta_pref + np.pi) % (2 * np.pi)
   signed_dist_2 = np.arctan2(np.sin(theta - theta_antipref), np.cos(theta - theta_antipref))
   # Determine effective tuning width for the second peak based on signed distance
   sigma_2_eff = np.where(signed_dist_2 < 0, tuning_width_2_left, tuning_width_2_right)
   # Calculate response for the second peak
   peak_2_response = amplitude_2 * np.exp(-0.5 * (np.abs(signed_dist_2) / sigma_2_eff) **
2)
   return baseline + peak_1_response + peak_2_response
~~~

### 4. Output for parameter estimator (Generation 1)

~~~
import numpy as np
def parameter_estimator(theta, spike_counts):
    “““
    Estimates parameters for neuron_model_v3. This estimator is adapted from parameter_estimator_v2,
    but adds estimation for the asymmetric tuning widths. “““
    n_bins = 50
    bin_idx = ((theta * n_bins) / (2 * np.pi)).astype(np.int32)
    bin_idx = np.clip(bin_idx, 0, n_bins - 1)
    sums = np.bincount(bin_idx, weights=spike_counts, minlength=n_bins)
    counts = np.bincount(bin_idx, minlength=n_bins)
    def gaussian_kernel(sig: int) -> np.ndarray:
        x = np.arange(-int(3 * sig), int(3 * sig) + 1)
        k = np.exp(-0.5 * (x / sig) ** 2)
        return k / np.sum(k)
    k = gaussian_kernel(2)
    pad = len(k) // 2
    sums_padded = np.pad(sums, (pad, pad), mode=‘wrap’)
    counts_padded = np.pad(counts, (pad, pad), mode=‘wrap’)
    num_conv = np.convolve(sums_padded, k, mode=‘valid’)
    den_conv = np.convolve(counts_padded, k, mode=‘valid’)
    tuning_curve = num_conv / (den_conv + 1e-8)

    pref_idx = np.argmax(tuning_curve)
    theta_pref = pref_idx * (2 * np.pi / n_bins)
    baseline = np.min(tuning_curve)
    amplitude_1 = np.max(tuning_curve) - baseline
    amplitude_2 = tuning_curve[(pref_idx + n_bins // 2) % n_bins] - baseline

    # Rough estimate of tuning widths - can be further refined
    tuning_width_1_left = np.pi / 4 # Initial guess
    tuning_width_1_right = np.pi / 4 # Initial guess
    tuning_width_2_left = np.pi / 4
    tuning_width_2_right = np.pi / 4

    return np.array([theta_pref, baseline, amplitude_1, tuning_width_1_left,
 tuning_width_1_right, amplitude_2, tuning_width_2_left, tuning_width_2_right])
~~~

### 5. Prompt for JAX Translator (Generation 1)

~~~
Convert the following function to a JAX-compatible function.
Include all necessary imports, and ensure that the function is compatible with JAX
transformations like ‵jax.jit‵, ‵jax.grad‵, and ‵jax.vmap‵.
Do not include any text other than the code.
Here is the code to translate:
import numpy as np
def neuron_model(theta, theta_pref=0.0, baseline=0.0, amplitude_1=1.0,
tuning_width_1_left=1.0, tuning_width_1_right=1.0, amplitude_2=0.0,
tuning_width_2_left=1.0, tuning_width_2_right=1.0):
”””
          parent_model_1: simple_gaussian_tuning_model
          parent_model_2: double_gaussian_tuning_model
Equation for this model:
R(theta) = baseline +
         amplitude_1 * exp(-0.5 * (abs(signed_dist_1) / sigma_1_eff)^2) +
         amplitude_2 * exp(-0.5 * (abs(signed_dist_2) / sigma_2_eff)^2)
where:
signed_dist_1 = arctan2(sin(theta - theta_pref), cos(theta - theta_pref))
sigma_1_eff = tuning_width_1_left if signed_dist_1 < 0 else tuning_width_1_right
signed_dist_2 = arctan2(sin(theta - (theta_pref + pi)), cos(theta - (theta_pref + pi)))
sigma_2_eff = tuning_width_2_left if signed_dist_2 < 0 else tuning_width_2_right
This model improves upon previous models by providing greater flexibility in tuning curve shapes, particularly by
     allowing for asymmetric Gaussian peaks and independent widths for the primary and secondary peaks.
The ‘simple_gaussian_tuning_model’ (neuron_model_v1) often failed to capture neurons with bi-directional tuning,
resulting in high loss for cells like Cell 469, Cell 683, Cell 67, and Cell 501, where it only fits one peak and
a flat baseline for the rest of the circle (e.g., green line for Cell 469 missing the peak around 4.5 rad).
The ‘double_gaussian_tuning_model’ (neuron_model_v2) was a significant improvement, introducing a second peak at
the anti-preferred direction. This dramatically reduced loss by fitting the second peak observed in the data mean
     (blue line) for many cells (e.g., comparing red vs green lines for Cell 358, Cell 469, Cell 683, Cell 915).
Despite this, the ‘double_gaussian_tuning_model’ assumed a shared and symmetric tuning_width for both peaks, which
        is often not true in biological data.
This model addresses the limitations of the ‘double_gaussian_tuning_model’ by:
1. Allowing the preferred (amplitude_1) and anti-preferred (amplitude_2) peaks to have *independent* widths.
   Even if ‵tuning_width_1_left‵ and ‵tuning_width_1_right‵ are equal (for symmetry), and similarly for
  the second peak, this model permits ‵tuning_width_1_left‵ (or ‵tuning_width_1_right‵) to be different from
‵tuning_width_2_left‵ (or ‵tuning_width_2_right‵). This enhanced flexibility can be beneficial where the
  main peak is broader or sharper than the anti-preferred peak, as can be visually suggested in Cell 67
  or Cell 469 where the peaks visually appear to have different extents.
2. Introducing *asymmetry* to each Gaussian peak, using separate width parameters for the left and right
     sides of each peak (e.g., tuning_width_1_left and tuning_width_1_right for the first peak). This is a crucial
feature for fitting real neural tuning curves, which are rarely perfectly symmetric Gaussians. For example,
in Cell 469, the main peak (around 1.5 rad) of the ‘Mean’ (blue curve) appears to have a steeper rise and a
     shallower decay, a characteristic not well-captured by the symmetric red ‘v2’ curve. Similarly, in Cell 501,
     the peak around 4 rad might benefit from asymmetric fitting to better match the blue mean curve.
    By modeling this asymmetry, this model can more precisely track the observed firing rate profiles, leading to
    a lower overall loss by minimizing systematic deviations between the model and the data’s true shape.
Args:
   theta (np.ndarray): The angle in radians for which to compute the neuron’s response.
   theta_pref (float): The preferred direction of the neuron for the primary peak.
   baseline (float): The baseline (minimum) firing rate of the neuron.
   amplitude_1 (float): The maximum firing rate above baseline for the primary peak.
tuning_width_1_left (float): The tuning width for the ‘left’ side (counter-clockwise) of the primary peak.
   tuning_width_1_right (float): The tuning width for the ‘right’ side (clockwise) of the primary peak.
   amplitude_2 (float): The maximum firing rate above baseline for the secondary peak (at theta_pref + pi).
   tuning_width_2_left (float): The tuning width for the ‘left’ side (counter-clockwise) of the secondary peak.
  tuning_width_2_right (float): The tuning width for the ‘right’ side (clockwise) of the secondary peak.
Returns:
    np.ndarray: The firing rate of the neuron at angle theta.
“““
 # Clip parameters to biologically plausible ranges
 theta_pref = np.clip(theta_pref, 0, 2 * np.pi)
 baseline = np.clip(baseline, 0, None) # Firing rate cannot be negative
 amplitude_1 = np.clip(amplitude_1, 0, None)
 amplitude_2 = np.clip(amplitude_2, 0, None)
 # Tuning widths must be positive and non-zero to avoid division by zero
 tuning_width_1_left = np.clip(tuning_width_1_left, 0.01, None)
 tuning_width_1_right = np.clip(tuning_width_1_right, 0.01, None)
 tuning_width_2_left = np.clip(tuning_width_2_left, 0.01, None)
 tuning_width_2_right = np.clip(tuning_width_2_right, 0.01, None)
 # Calculate signed circular distance for the first peak
 signed_dist_1 = np.arctan2(np.sin(theta - theta_pref), np.cos(theta - theta_pref))
 # Determine effective tuning width for the first peak based on signed distance
sigma_1_eff = np.where(signed_dist_1 < 0, tuning_width_1_left, tuning_width_1_right)
 # Calculate response for the first peak
peak_1_response = amplitude_1 * np.exp(-0.5 * (np.abs(signed_dist_1) / sigma_1_eff) ** 
2)
 # Calculate signed circular distance for the second peak (anti-preferred direction)
theta_antipref = (theta_pref + np.pi) % (2 * np.pi)
signed_dist_2 = np.arctan2(np.sin(theta - theta_antipref), np.cos(theta - theta_antipref))
 # Determine effective tuning width for the second peak based on signed distance
sigma_2_eff = np.where(signed_dist_2 < 0, tuning_width_2_left, tuning_width_2_right)
 # Calculate response for the second peak
peak_2_response = amplitude_2 * np.exp(-0.5 * (np.abs(signed_dist_2) / sigma_2_eff) ** 
2)
return baseline + peak_1_response + peak_2_response
~~~

### 6. Output for JAX Translator (Generation 1)

~~~
import jax
import jax.numpy as jnp
def neuron_model(theta, theta_pref=0.0, baseline=0.0, amplitude_1=1.0,
tuning_width_1_left=1.0, tuning_width_1_right=1.0, amplitude_2=0.0,
tuning_width_2_left=1.0, tuning_width_2_right=1.0):
     theta_pref = jnp.clip(theta_pref, 0, 2 * jnp.pi) baseline = jnp.clip(baseline, 0, None)
     amplitude_1 = jnp.clip(amplitude_1, 0, None)
     amplitude_2 = jnp.clip(amplitude_2, 0, None)
     tuning_width_1_left = jnp.clip(tuning_width_1_left, 0.01, None)
     tuning_width_1_right = jnp.clip(tuning_width_1_right, 0.01, None)
     tuning_width_2_left = jnp.clip(tuning_width_2_left, 0.01, None)
     tuning_width_2_right = jnp.clip(tuning_width_2_right, 0.01, None)
     signed_dist_1 = jnp.arctan2(jnp.sin(theta - theta_pref), jnp.cos(theta - theta_pref))
     sigma_1_eff = jnp.where(signed_dist_1 < 0, tuning_width_1_left, tuning_width_1_right)
     peak_1_response = amplitude_1 * jnp.exp(-0.5 * (jnp.abs(signed_dist_1) / sigma_1_eff) 
** 2)
    theta_antipref = (theta_pref + jnp.pi) % (2 * jnp.pi)
    signed_dist_2 = jnp.arctan2(jnp.sin(theta - theta_antipref), jnp.cos(theta - 
theta_antipref))
    sigma_2_eff = jnp.where(signed_dist_2 < 0, tuning_width_2_left, tuning_width_2_right)
    peak_2_response = amplitude_2 * jnp.exp(-0.5 * (jnp.abs(signed_dist_2) / sigma_2_eff) ** 2)
   return baseline + peak_1_response + peak_2_response
~~~

### 7. Prompt that generated winning neuron model

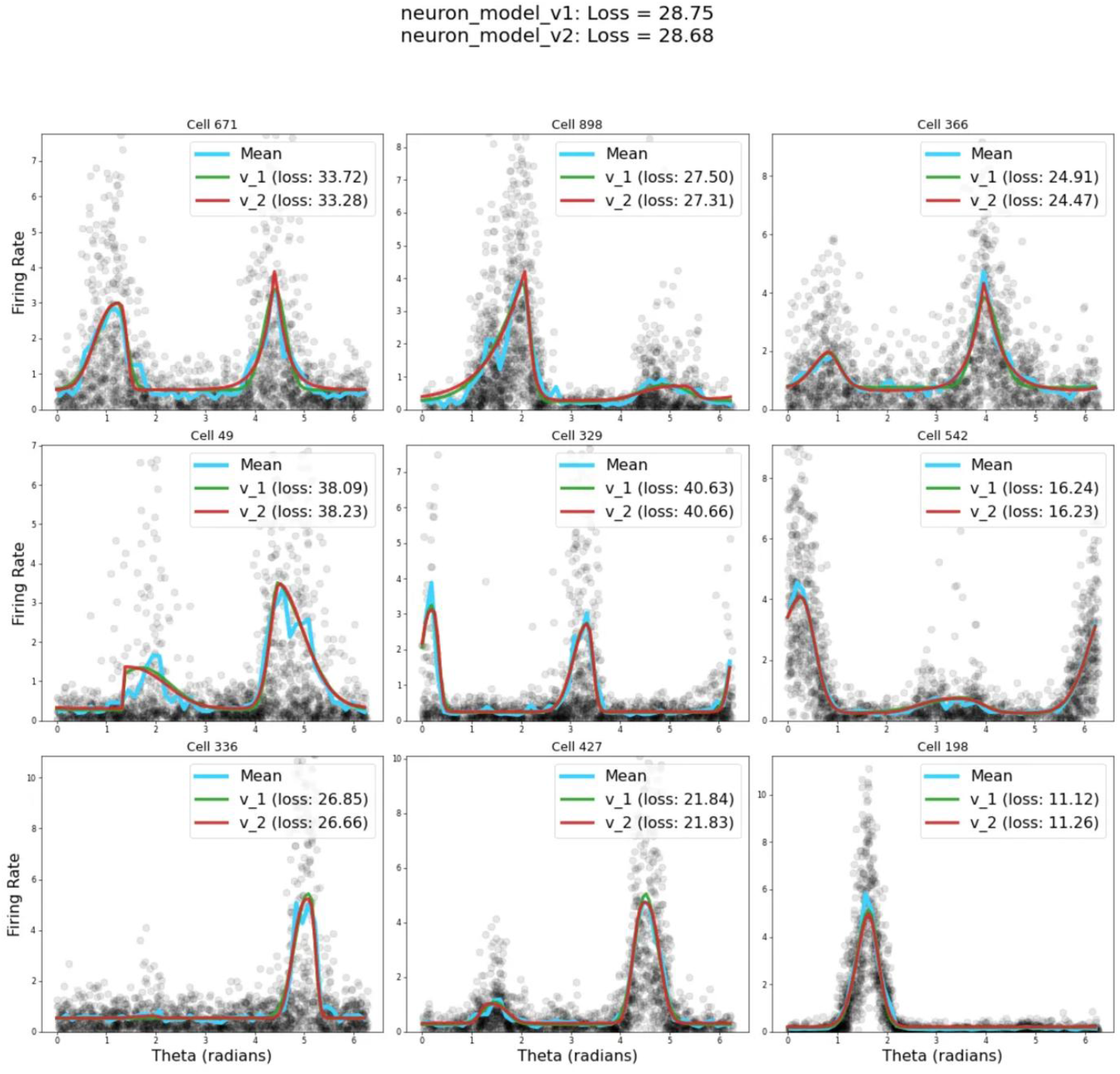

~~~
You are an AI scientist. The programs below are biological models of neurons. The models are sorted from highest to lowest loss.
Your task is to create a new neuron model, neuron_model_v3, that has a lower loss than the models below.
*Analyze* the progression of the models, *generalize* the improvements, and *create* a new model that is better than *all* previous models.
Use the models below as a *template* to create a new model.
Which features in the models below correlate with lower loss? Find these features and 
*extrapolate* them.
Focus on *exploiting* the strengths of the existing models and *eliminating* their weaknesses or *redundancies*.
Are the parameter ranges correct? If not, adjust them to be more appropriate.
You will be *penalized* for complexity, so make the new model as *simple* as possible while still being better than the previous models.
**Image Analysis Instructions:**
Attached is a scatter plot of the neuron models’ performance on top of raw neural data. The binned mean is plotted in **sky-blue**, ‵neuron_model_v1‵ is plotted in **green**, and ‵neuron_model_v2‵ is plotted in **red**.
Analyse the models’ fits to the data in the image below. Identify systematic weaknesses of the models by observing patterns across multiple cell plots. For instance, consider:
**Model Comparisons:** Which models are better for each cell? That is to say, which models track the blue curve better? Which features of the models are responsible for improving the fit?
**Model Fit:** How well do the models fit the binned data mean? Look for places where even the models (**red** curve for best model, **green** for second best model) deviate most from the binned data mean (**blue** curve). This is where the models are weakest, and where you should focus your improvements.
**Model Shape:** Do the models’ shapes (e.g., peak sharpness, width, skewness, amplitude, etc.) align with the binned data mean (**blue**) and raw data scatter points (**black**)? If not, how do they differ? How can you change the model to better match the data shape?
**Parameter Flexibility:** Are there free parameters that could be introduced or modified to better capture the observed response profiles? Utilize your analysis of the shortcomings of the current models’ shapes and add free parameters or modify existing ones to address these issues.
Use this analysis to inform the design of a new neuron model, ‵neuron_model_v3‵, that improves upon the previous models.
Include your analysis of the image in the docstring of your new model. Point to specific subplots in the image that illustrate the *strengths* and *weaknesses* of the parent models. Explain how you plan to **fix** the weaknesses of the parent models.
**Code Generation Guidelines:**
* Import any packages you use.
* Do not include any text other than the code.
* Ensure all free parameters are numeric, not strings.
* At the beginning of the code, clip the free parameters to a biologically plausible range, e.g., ‵theta_pref = np.clip(theta_pref, 0, 2 * jnp.pi)‵.
**Docstring Guidelines:**
* Begin by listing the parent models and give them a name that describes their key features, e.g., ‵parent_model_1: simple_exponential_decay-model‵, ‵parent_model_2: double_exponential_decay_model‵. Never refer to the models as ‵neuron_model_v1‵, ‵neuron_model_v2‵, etc. Instead, refer to them as ‵parent_models‵ or their descriptive names (e.g. ‵simple_exponential_decay_model‵).
* Do not refer to the current model as ‵neuron_model_v3‵. Instead, refer to it as “this model”.
* Provide a simple equation for the model, including all free parameters.
* Include a brief description of how the model improves upon the previous models, citing specific features or changes that lead to lower loss.
~~~

**`loss of model 1: 28.75`**

~~~
import numpy as np
import jax.numpy as jnp
 def neuron_model_v1(theta, theta_pref=0.0, baseline=0.0, 
                      amplitude_1=1.0, tuning_width_1_left=1.0, tuning_width_1_right=1.0,
skewness_1=0.0, 
                      amplitude_2=0.0, tuning_width_2_left=1.0, tuning_width_2_right=1.0,
skewness_2=0.0): “““
Parent Model 1: Asymmetric Double Gaussian Tuning Curve (neuron_model_v2)
       Equation: ‵f(θ) = B + A_1_ * exp(-0.5 * (Δθ_1_ / σ_1_(Δθ_1_≥0 ? R : L))^2^) + A_2_ * exp(-0.5 * (Δθ_2_ / σ_2_(Δθ_2_≥0 ? R : L))^2^)‵, where Δθ is circular distance, B is baseline, A is amplitude, and σ represents the tuning width to the left (L) or right (R).
       Strengths: Captures asymmetry and two preferred directions. Asymmetry fixes some of the fit issues evident across plots, especially in cell 1035.
Parent Model 2: Flexible Double Gaussian Tuning Curve (neuron_model_v1) 
      Equation:
       ‵firing_rate = baseline + amplitude_1 * exp(-0.5 * (circ_dist(theta, theta_pref) / tuning_width_1)^2) + amplitude_2 * exp(-0.5 * (circ_dist(theta, (theta_pref + pi + angle_offset_2) % (2 * pi)) / tuning_width_2)^2)‵
      Strengths: allows the 2nd peak to be offset, fixing the fit issues for Cell 193.
This Model: Skewed Asymmetric Double Gaussian Tuning Curve
   Equation:
        ‵f(θ) = B + A_1_ * exp(-0.5 * (Δθ_1_ / s_1_(Δθ_1_≥0 ? R : L))^2^ + skewness_1*Δθ_1_) + A_2_ * exp(-0.5 * (Δθ_2_ / s_2_(Δθ_2_≥0 ? R : L))^2^ + skewness_2*Δθ_2_)‵
       Where:
       *A_1_, A_2_: Amplitudes of peak 1 and peak 2, respectively.
       *Δθ_1_, Δθ_2_: Angular distances from the data angle ‵theta‵ to the centre of peak 1 and peak 2, respectively
      *s_1_(Δθ_1_≥0 ? R : L) and s_2_(Δθ_2_≥0 ? R : L): tuning widths (standard deviations) of peak 1 and peak 2, where separate parameters are defined to the left and right.
     *skewness_1 and skewness_2: additional weighting, where separate parameters can skew each of the peaks of each Gaussian curve.
     Improvements:
          Building upon the asymmetric tuning width, a skewness parameter is added to each Gaussian that models the shape of each response more closely.
Specifically:
           *Addresses shape issues for peaks such as in Cells 1035 and 254
           *Increases robustness of the model when 1st and 2nd tuning responses aren’t symmetrical or equal
   “““
      # Parameter clipping to biologically plausible ranges and to avoid division by zero/singularities
      theta_pref = np.clip(theta_pref, 0, 2 * jnp.pi)
      baseline = np.clip(baseline, 0, None) # Must be non-negative
      amplitude_1 = np.clip(amplitude_1, 0, None) # Must be non-negative
      amplitude_2 = np.clip(amplitude_2, 0, None) # Must be non-negative
     # Tuning widths must be positive to ensure Gaussian shape. Add small epsilon to prevent zero.
     tuning_width_1_left = np.clip(tuning_width_1_left, 1e-3, None)
     tuning_width_1_right = np.clip(tuning_width_1_right, 1e-3, None)
     tuning_width_2_left = np.clip(tuning_width_2_left, 1e-3, None)
     tuning_width_2_right = np.clip(tuning_width_2_right, 1e-3, None)
    skewness_1 = np.clip(skewness_1, -1, 1)
    skewness_2 = np.clip(skewness_2, -1, 1)
    # Helper function to compute signed circular difference
    # This correctly computes difference in [-pi, pi]
   circ_signed_diff = lambda t1, t2: jnp.arctan2(jnp.sin(t1 - t2), jnp.cos(t1 - t2))
    # Calculate response for the first peak (around theta_pref) delta_theta_1 = circ_signed_diff(theta, theta_pref)
    # Select the appropriate tuning width based on the sign of the difference
   current_tuning_width_1 = jnp.where(delta_theta_1 >= 0, 
tuning_width_1_right, tuning_width_1_left)
   response_peak_1 = amplitude_1 * jnp.exp(-0.5 * (delta_theta_1 / current_tuning_width_1)**2 + skewness_1*delta_theta_1)
    # Calculate response for the second peak (around theta_pref + pi)
   theta_pref_2 = (theta_pref + jnp.pi) % (2 * jnp.pi)
   delta_theta_2 = circ_signed_diff(theta, theta_pref_2)
    # Select the appropriate tuning width based on the sign of the difference
current_tuning_width_2 = jnp.where(delta_theta_2 >= 0, tuning_width_2_right, tuning_width_2_left)
    response_peak_2 = amplitude_2 * jnp.exp(-0.5 * (delta_theta_2 / current_tuning_width_2)**2 + skewness_2*delta_theta_2)
   
    # Combine baseline and two peak responses
   return baseline + response_peak_1 + response_peak_2
~~~

**`loss of model 2: 28.68`**

~~~
import numpy as np
def neuron_model_v2(theta, theta_pref=0.0, baseline=0.0, amplitude_1=1.0, amplitude_2=0.0,
                    tuning_width_1_left=1.0,
                    tuning_width_1_right=1.0,
                    tuning_width_2_left=1.0,
                    tuning_width_2_right=1.0,
                    exponent_1=2.0, exponent_2=2.0):
“““
   parent_model_1: unimodal_gaussian_model (neuron_model_v1)
Equation: ‵R(theta) = B + A * exp(-0.5 * (d(theta, P_1) / W)^2‵
   parent_model_2: bimodal_asymmetric_gaussian_model (neuron_model_v2)
   Equation: ‵R(theta) = B + A_1 * exp(-0.5 * (d_s(theta, P_1) / W_{1,eff})^2) + A_2 * 
exp(-0.5 * (d_s(theta, P_2) / W_{2,  eff})^2)‵
   where ‵P_2 = (P_1 + pi) % (2*pi)‵.
   This model: asymmetric_double_generalized_gaussian_tuning_model
   Equation: ‵R(theta) = B + A_1 * exp(-0.5 * (abs(d_s(theta, P_1)) / W_{1,eff})^E_1) + A_2 * exp(-0.5 * (abs(d_s(theta, P_2)) / W_{2,eff})^E_2)‵
   
  Where:
  - ‵B‵ is the baseline firing rate.
  - ‵A_1‵, ‵A_2‵ are the amplitudes of the first and second peaks.
  - ‵P_1‵ is the preferred direction of the first peak (‵theta_pref‵).
  - ‵P_2 = (P_1 + pi) % (2*pi)‵ is the preferred direction of the second (opposite) peak.
  - ‵d_s(theta, P)‵ denotes the signed circular distance: ‵arctan2(sin(theta - P), cos(theta - P))‵.
  - ‵W_{i,eff}‵ is the effective tuning width for peak ‵i‵, chosen as ‵W_{i,right}‵ if ‵d_s(theta, P_i) >= 0‵ and ‵W_{i,left}‵ if ‵d_s(theta, P_i) < 0‵.
  - ‵E_1‵ and ‵E_2‵ are the exponents that control the sharpness of the peaks. ‵E‵ close to zero results in broad plateaus at the peak, while a larger ‵E‵ gives increasingly sharp peaks.
This model builds upon the strengths of ‵bimodal_asymmetric_gaussian_model‵ and addresses remaining weaknesses observed in the plots:
  1. **Generalized Gaussian Shape:** The ‵bimodal_asymmetric_gaussian_model‵ assumes a Gaussian shape (exponent fixed at 2). While the previous model allowed asymmetric widths, it still might not capture the sharpness or flatness of the peaks effectively. For example, some neurons might have very sharp peaks (e.g., Cell 471 around 5 radians) which are poorly fit by a gaussian-like shape, even with asymmetrical widths. Other cells might have flatter, plateau-like regions near their peak. To address this limitation, this model introduces exponents (‵exponent_1‵ and ‵exponent_2‵) for both peaks, allowing the tuning curve to be more or less sharply peaked than a Gaussian. This modification enhances the flexibility in fitting the peak shape, adapting to various observed firing rate profiles.
  2. **Independent Exponents**: ‵exponent_1‵ and ‵exponent_2‵ independently control the peak sharpness of the main and anti-preferred direction. ‵Cell 672‵ demonstrates that a neuron’s preferred and anti-preferred directions need not have tuning curves of the same shape, therefore it’s important that peak shape (controlled by the exponent) can be independently adjusted.
  3. **Absolute Signed Distance**: In order to guarantee proper peak shape the *absolute value* of the signed distance measure, i.e. ‵abs(d_s(theta, P_1))‵, must be used as an argument in the equation, because a negative value to the power of exponent ‵E‵ produces complex numbers which should not be returned by this real valued function. The downside of using an absolute value is the possibility of some minor discontinuities in the deriviatives, but in this case visual peak quality is given preference over differentiability.
Args:
    theta (np.ndarray): The angle in radians.
    theta_pref (float): Preferred direction of the main peak (first peak).
    baseline (float): Baseline firing rate.
    amplitude_1 (float): Maximum firing rate above baseline for the first peak.
    amplitude_2 (float): Maximum firing rate above baseline for the second (opposite) 
peak.
    tuning_width_1_left (float): Width parameter for the “left” side (negative signed distance) of the first peak.
    tuning_width_1_right (float): Width parameter for the “right” side (positive signed distance) of the first peak.
    tuning_width_2_left (float): Width parameter for the “left” side (negative signed distance) of the second peak.
    tuning_width_2_right (float): Width parameter for the “right” side (positive signed distance) of the second peak.
    exponent_1 (float): Exponent that controls the sharpness of the first peak.
    exponent_2 (float): Exponent that controls the sharpness of the second peak.
Returns:
    np.ndarray: The firing rate of the neuron at angle theta.
“““
    # Clip parameters to biologically plausible ranges
    theta_pref = np.clip(theta_pref, 0, 2 * np.pi)
    baseline = np.clip(baseline, 0, None)
    amplitude_1 = np.clip(amplitude_1, 0, None)
    amplitude_2 = np.clip(amplitude_2, 0, None)
    tuning_width_1_left = np.clip(tuning_width_1_left, 0.01, None)
    tuning_width_1_right = np.clip(tuning_width_1_right, 0.01, None)
    tuning_width_2_left = np.clip(tuning_width_2_left, 0.01, None)
    tuning_width_2_right = np.clip(tuning_width_2_right, 0.01, None)
    exponent_1 = np.clip(exponent_1, 0.1, 10) # Restrict to avoid numerical instability
    exponent_2 = np.clip(exponent_2, 0.1, 10) # Restrict to avoid numerical instability 
    # Calculate signed circular distance
      circ_signed_dist = lambda theta_val, theta_p: np.arctan2(np.sin(theta_val - theta_p), np.cos(theta_val - theta_p))
    
   # --- Peak 1 ---
  dist_1_signed = circ_signed_dist(theta, theta_pref)
   # Select tuning width based on the sign of the distance (left/right of peak)
   tuning_width_1_eff = np.where(dist_1_signed >= 0, tuning_width_1_right, tuning_width_1_left)
   # Calculate response for Peak 1, use abs to guarantee function remains real
   peak_1_response = amplitude_1 * np.exp(-0.5 * (np.abs(dist_1_signed) / 
tuning_width_1_eff) ** exponent_1)
   # --- Peak 2 (at opposite direction) --- 
  theta_pref_2 = (theta_pref + np.pi) % (2 * np.pi) 
  dist_2_signed = circ_signed_dist(theta, theta_pref_2)
  # Select tuning width based on the sign of the distance (left/right of peak)
  tuning_width_2_eff = np.where(dist_2_signed >= 0, tuning_width_2_right, 
tuning_width_2_left)
  # Calculate response for Peak 2, use abs to guarantee function remains real
  peak_2_response = amplitude_2 * np.exp(-0.5 * (np.abs(dist_2_signed) / tuning_width_2_eff) ** exponent_2)
  # Total response
  return baseline + peak_1_response + peak_2_response
~~~

### 8. Output for winning neuron model

~~~
import numpy as np
import jax.numpy as jnp
def neuron_model(theta, theta_pref=0.0, baseline=0.0, amplitude_1=1.0, amplitude_2=0.0,
                     tuning_width_1_left=1.0,tuning_width_1_right=1.0,
                     tuning_width_2_left=1.0, tuning_width_2_right=1.0,
                     exponent_1=2.0, exponent_2=2.0, angle_offset_2=0.0):
“““
Parent Model 1: Skewed Asymmetric Double Gaussian Tuning Curve (neuron_model_v1)
       Equation:
       ‵f(θ) = B + A_1_ * exp(-0.5 * (Δθ_1_ / s_1_(Δθ_1_≥0 ? R : L))^2^ + skewness_1*Δθ_1_) + A_2_ * 
exp(-0.5 * (Δθ_2_ / s_2_(Δθ_2_≥0 ? R : L))^2^ + skewness_2*Δθ_2_)‵
Parent Model 2: asymmetric_double_generalized_gaussian_tuning_model (neuron_model_v2)
     Equation: ‵R(theta) = B + A_1 * exp(-0.5 * (abs(d_s(theta, P_1)) / W_{1,eff})^E_1) 
+ A_2 * exp(-0.5 * (abs(d_s(theta, P_2) / W_{2,eff}))^E_2)‵
    Where ‵P_2 = (P_1 + pi) % (2*pi)‵.
This Model: Offset Asymmetric Double Generalized Gaussian Tuning Curve
    Equation:
    ‵R(theta) = B + A_1 * exp(-0.5 * (abs(d_s(theta, P_1)) / W_{1,eff})^E_1) + A_2 * exp(-0.5 * (abs(d_s(theta, P_2) / W_{2,eff}))^E_2)‵
Where:
   * B: Baseline firing rate
   *A_1, A_2: Amplitudes of peak 1 and peak 2
   *P_1: Preferred direction of the first peak (‵theta_pref‵)
   *P_2: Preferred direction of the second peak, where ‵P_2 = (P_1 + pi + angle_offset_2) % (2 * pi)‵
   *d_s(theta, P): Signed circular distance between theta and P
   *W_{i,eff}: Effective tuning width for peak i (left or right, depending on the sign of d_s)
  *E_1, E_2: Exponents controlling the peak sharpness
Improvements:
    Building upon the asymmetric and generalized gaussian model, this version adds an 
offset parameter to the second peak (angle_offset_2).
    The introduction of the ‵angle_offset_2‵ parameter enables the model to account for 
scenarios where the second peak is not precisely opposite the primary peak.
   This is beneficial, for example, in cell 671 where the secondary peak is offset to the left (i.e., smaller angle) relative to the opposite (π radians away) side of the 
primary peak.
The main reason to choose ‵angle_offset_2‵ for improvement is due to its low complexity and the ready-to-use building blocks of the two parent models, thereby improving 
the parent model with a singular cost-effective addition.
    “““
    # Clip parameters to biologically plausible ranges
    theta_pref = np.clip(theta_pref, 0, 2 * jnp.pi)
    baseline = np.clip(baseline, 0, None)
    amplitude_1 = np.clip(amplitude_1, 0, None)
    amplitude_2 = np.clip(amplitude_2, 0, None)
    tuning_width_1_left = np.clip(tuning_width_1_left, 0.01, None)
    tuning_width_1_right = np.clip(tuning_width_1_right, 0.01, None)
    tuning_width_2_left = np.clip(tuning_width_2_left, 0.01, None)
    tuning_width_2_right = np.clip(tuning_width_2_right, 0.01, None)
    exponent_1 = np.clip(exponent_1, 0.1, 10)
    exponent_2 = np.clip(exponent_2, 0.1, 10)
    angle_offset_2 = np.clip(angle_offset_2, -jnp.pi, jnp.pi)
    # Calculate signed circular distance
      circ_signed_dist = lambda theta_val, theta_p: jnp.arctan2(jnp.sin(theta_val - theta_p), np.cos(theta_val - theta_p))
    # --- Peak 1 ---
    dist_1_signed = circ_signed_dist(theta, theta_pref)
    # Select tuning width based on the sign of the distance (left/right of peak)
    tuning_width_1_eff = jnp.where(dist_1_signed >= 0, tuning_width_1_right, 
tuning_width_1_left)
    # Calculate response for Peak 1, use abs to guarantee function remains real
    peak_1_response = amplitude_1 * jnp.exp(-0.5 * (jnp.abs(dist_1_signed) / 
tuning_width_1_eff) ** exponent_1)
    # --- Peak 2 (at opposite direction, with offset) ---
    theta_pref_2 = (theta_pref + jnp.pi + angle_offset_2) % (2 * jnp.pi)
    dist_2_signed = circ_signed_dist(theta, theta_pref_2)
   # Select tuning width based on the sign of the distance (left/right of peak) tuning_width_2_eff = jnp.where(dist_2_signed >= 0, tuning_width_2_right,
tuning_width_2_left)
   
   # Calculate response for Peak 2, use abs to guarantee function remains real
peak_2_response = amplitude_2 * jnp.exp(-0.5 * (jnp.abs(dist_2_signed) / tuning_width_2_eff) ** exponent_2)
   # Total response
   return baseline + peak_1_response + peak_2_response
~~~

### 9. Parameter Estimator of Winning Model

~~~
import jax.numpy as jnp
def parameter_estimator(theta, spike_counts):
    n_bins = 75
    bin_idx = ((theta * n_bins) / (2 * np.pi)).astype(np.int32)
    bin_idx = np.clip(bin_idx, 0, n_bins - 1)
    sums = np.bincount(bin_idx, weights=spike_counts, minlength=n_bins)
    counts = np.bincount(bin_idx, minlength=n_bins)
    k = np.exp(-0.5 * (np.arange(-5, 6)) ** 2)
    k /= np.sum(k)
    sums_padded = np.pad(sums, (5, 5), mode=‘wrap’)
    counts_padded = np.pad(counts, (5, 5), mode=‘wrap’)
tuning_curve = np.convolve(sums_padded, k, mode=‘valid’) / (np.convolve(counts_padded, 
k, mode=‘valid’) + 1e-8)
   baseline = np.min(tuning_curve)
   peak_1_idx = np.argmax(tuning_curve)
   theta_pref = peak_1_idx * (2 * np.pi / n_bins)
   amplitude_1 = tuning_curve[peak_1_idx] – baseline
   anti_pref_idx = (peak_1_idx + n_bins // 2) % n_bins
   amplitude_2 = max(0, tuning_curve[anti_pref_idx] - baseline)
   theta_pref_2_est = anti_pref_idx * (2 * np.pi / n_bins)
   angle_offset_2 = theta_pref_2_est - (theta_pref + np.pi) % (2 * np.pi)
tuning_width_1_left = tuning_width_1_right = tuning_width_2_left = tuning_width_2_right = np.pi / 8
   exponent_1 = exponent_2 = 2.0
   return np.array([theta_pref, baseline, amplitude_1, amplitude_2, 
tuning_width_1_left, tuning_width_1_right, tuning_width_2_left, tuning_width_2_right, exponent_1, exponent_2, 
angle_offset_2])
~~~

## Supplementary Text 3: Equations for winner’s family tree

This appendix lists the LLM-generated equations in the winning model’s “family tree” for one run (Figure 2h). All models except B12 consist of different combinations of the features that eventually appear in the winning model H27. Model B12 adds a linear term to the exponential argument, which effectively shifts the Gaussian mean separately to the left and right of the tuning curve peaks; this feature was not selected by further evolution.

Of all 8 runs, 6 found the same solution as H27, for score 0.20. One run’s winner was the same as B35, with score of 0.19; the other was like H27 but with only one peak asymmetrical, for score 0.18.

### Graphical summary of equations fit in one example run

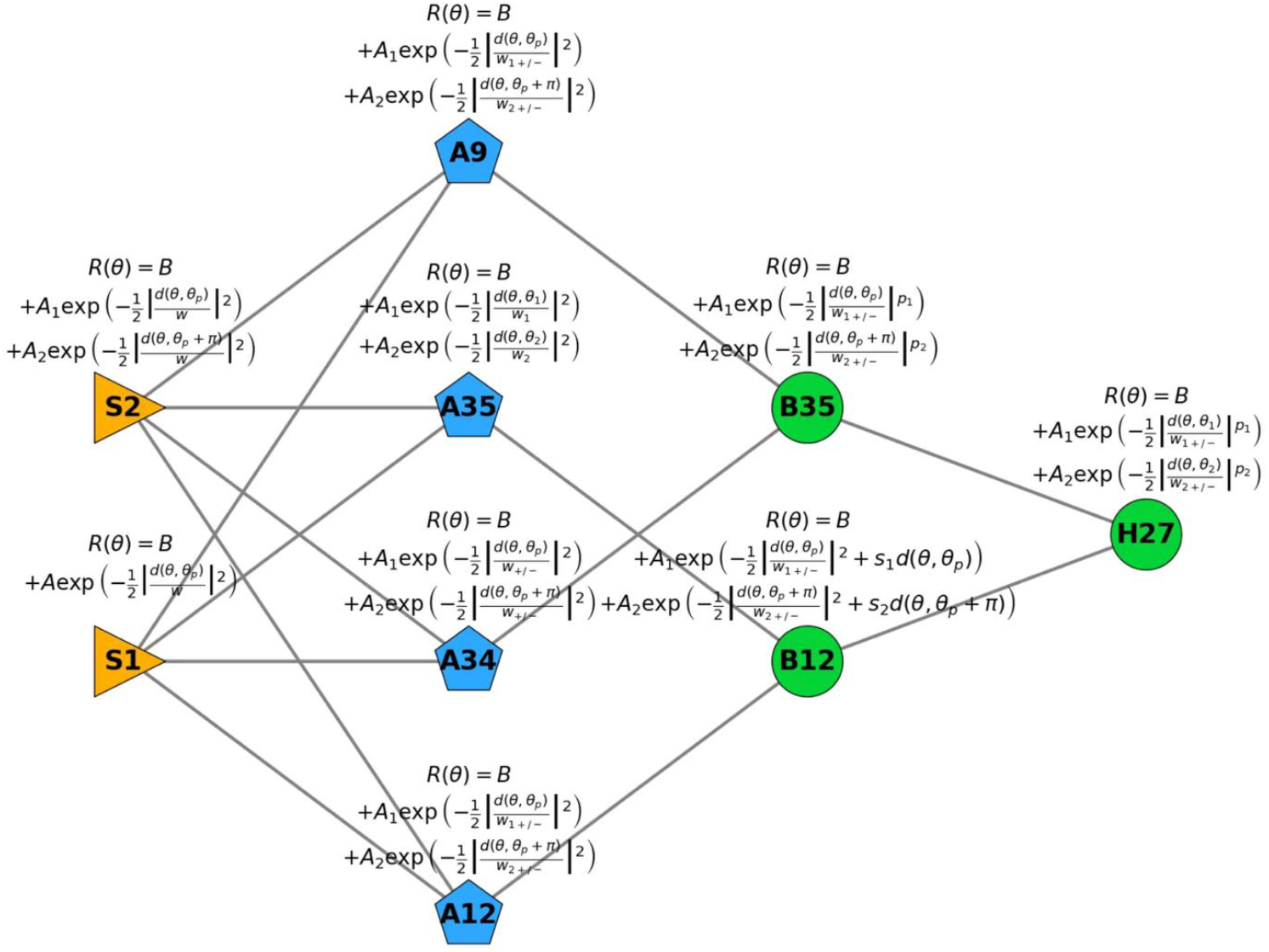

### Equation list

**Model S1, human-written seed model: single-Gaussian**

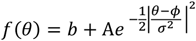

**Model S2, human-written seed model: double-Gaussian**

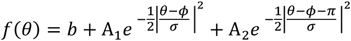

**Model A9, generation 1, Gemini 2.5: Asymmetric, different widths for different peaks**

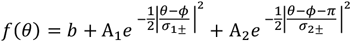

**Model A35, generation 1, Gemini 2.5: Non-antipodial, different symmetric widths for different peaks**

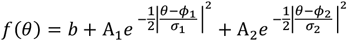

**Model A34, generation 1, Gemini 2.5: Asymmetric, same widths for different peaks**

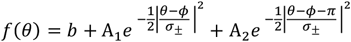

**Model A12, generation 1, Gemini 2.5: same as A9**

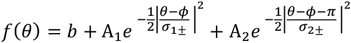

**Model B35, generation 2, Gemini 2.0: Non-Gaussian, asymmetric, different widths, but still antipodal**

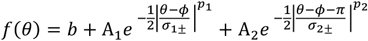

**Model B12, generation 2, Gemini 2.0: Asymmetric, different widths, linear term**

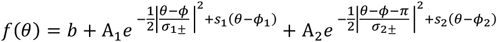

**Winner H27; generation 8, Gemini 2.0: Non-Gaussian, asymmetric, different widths, non-antipodal**

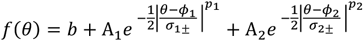

## References

1. Cover, T. M. Geometrical and Statistical Properties of Systems of Linear Inequalities with Applications in Pattern Recognition. IEEE Transactions on Electronic Computers EC-14, 326– 334 (1965).

2. Marr, D. A theory of cerebellar cortex. J.Physiol 202, 437–470 (1969).

3. Marr, D. A theory for cerebral neocortex. Proc.R.Soc.Lond B Biol.Sci. 176, 161–234 (1970).

4. Barlow, H. B. & Rosenblith, W. A. Possible principles underlying the transformation of sensory messages. in Sensory communication (MIT Press, Cambridge MA, 1961).

5. Attneave, F. Some informational aspects of visual perception. Psychol.Rev. 61, 183–193 (1954).

6. Chung, S. & Abbott, L. F. Neural population geometry: An approach for understanding biological and artificial neural networks. Current Opinion in Neurobiology 70, 137–144 (2021).

7. Fusi, S., Miller, E. K. & Rigotti, M. Why neurons mix: high dimensionality for higher cognition. Current Opinion in Neurobiology 37, 66–74 (2016).

8. DiCarlo, J. J., Zoccolan, D. & Rust, N. C. How does the brain solve visual object recognition? Neuron 73, 415–34 (2012).

9. Simoncelli, E. P. & Olshausen, B. A. Natural image statistics and neural representation. Annu Rev Neurosci 24, 1193–1216 (2001).

10. Bordelon, B. & Pehlevan, C. Population codes enable learning from few examples by shaping inductive bias. eLife 11, e78606 (2022).

11. Chou, C.-N. et al. Geometry Linked to Untangling Efficiency Reveals Structure and Computation in Neural Populations. 2024.02.26.582157 Preprint at 10.1101/2024.02.26.582157 (2025).

12. Kaufman, M. T. et al. The implications of categorical and category-free mixed selectivity on representational geometries. Current Opinion in Neurobiology 77, 102644 (2022).

13. Tang, W. et al. A hippocampal population code for rapid generalization. 2025.03.15.643456 Preprint at 10.1101/2025.03.15.643456 (2025).

14. Froudarakis, E. et al. Population code in mouse V1 facilitates readout of natural scenes through increased sparseness. Nat Neurosci 17, 851–857 (2014).

15. Stringer, C., Pachitariu, M., Steinmetz, N., Carandini, M. & Harris, K. D. High-dimensional geometry of population responses in visual cortex. Nature 571, 361–365 (2019).

16. Swindale, N. V. Orientation tuning curves: empirical description and estimation of parameters. Biol Cybern 78, 45–56 (1998).

17. Carandini, M. & Ferster, D. Membrane potential and firing rate in cat primary visual cortex. The Journal of neuroscience : the official journal of the Society for Neuroscience 20, 470–84 (2000).

18. Yamins, D. L. K. & DiCarlo, J. J. Using goal-driven deep learning models to understand sensory cortex. Nat Neurosci 19, 356–365 (2016).

19. Yamins, D. L. K. et al. Performance-optimized hierarchical models predict neural responses in higher visual cortex. Proceedings of the National Academy of Sciences 111, 8619–8624 (2014).

20. Cadena, S. A. et al. Deep convolutional models improve predictions of macaque V1 responses to natural images. PLOS Computational Biology 15, e1006897 (2019).

21. Kindel, W. F., Christensen, E. D. & Zylberberg, J. Using deep learning to probe the neural code for images in primary visual cortex. Journal of Vision 19, 29 (2019).

22. Turishcheva, P. et al. Retrospective for the dynamic sensorium competition for predicting large-scale mouse primary visual cortex activity from videos. in Proceedings of the 38th International Conference on Neural Information Processing Systems vol. 37 118907–118929 (Curran Associates Inc., Red Hook, NY, USA, 2024).

23. Wang, E. Y. et al. Foundation model of neural activity predicts response to new stimulus types. Nature 640, 470–477 (2025).

24. Lindsey, J. W. & Issa, E. B. Factorized visual representations in the primate visual system and deep neural networks. eLife 13, RP91685 (2024).

25. Li, B. M., Cornacchia, I. M., Rochefort, N. L. & Onken, A. V1T: large-scale mouse V1 response prediction using a Vision Transformer. Preprint at 10.48550/arXiv.2302.03023 (2023).

26. Ecker, A. S. et al. A rotation-equivariant convolutional neural network model of primary Tilbury et al. AI-derived tuning laws explain population geometry visual cortex. Preprint at 10.48550/arXiv.1809.10504 (2018).

27. Udrescu, S.-M. & Tegmark, M. AI Feynman: A physics-inspired method for symbolic regression. Science Advances 6, eaay2631 (2020).

28. Cranmer, M. Interpretable Machine Learning for Science with PySR and SymbolicRegression.jl. Preprint at 10.48550/arXiv.2305.01582 (2023).

29. Real, E., Liang, C., So, D. R. & Le, Q. V. AutoML-Zero: Evolving Machine Learning Algorithms From Scratch. Preprint at 10.48550/arXiv.2003.03384 (2020).

30. Correa, C. G., Griffiths, T. L. & Daw, N. D. Program-Based Strategy Induction for Reinforcement Learning. arXiv.org https://arxiv.org/abs/2402.16668v1 (2024).

31. Goodman, N. D., Tenenbaum, J. B., Feldman, J. & Griffiths, T. L. A Rational Analysis of Rule-Based Concept Learning. Cognitive Science 32, 108–154 (2008).

32. Schulze Buschoff, L. M., Akata, E., Bethge, M. & Schulz, E. Visual cognition in multimodal large language models. Nat Mach Intell 7, 96–106 (2025).

33. Mitchener, L. et al. Kosmos: An AI Scientist for Autonomous Discovery. Preprint at 10.48550/arXiv.2511.02824 (2025).

34. Gottweis, J. et al. Towards an AI co-scientist. Preprint at 10.48550/arXiv.2502.18864 (2025).

35. Lu, C. et al. The AI Scientist: Towards Fully Automated Open-Ended Scientific Discovery. Preprint at 10.48550/arXiv.2408.06292 (2024).

36. Li, V. R., Sun, J. & Wattenberg, M. Does visualization help AI understand data? Preprint at 10.48550/arXiv.2507.18022 (2025).

37. Yu, Z. & Lee, Y. J. How Multimodal LLMs Solve Image Tasks: A Lens on Visual Grounding, Task Reasoning, and Answer Decoding. Preprint at 10.48550/arXiv.2508.20279 (2025).

38. Geng, J., Chen, H., Arumugam, D. & Griffiths, T. L. Are Large Language Models Reliable AI Scientists? Assessing Reverse-Engineering of Black-Box Systems. Preprint at 10.48550/arXiv.2505.17968 (2025).

39. Luo, X. et al. Large language models surpass human experts in predicting neuroscience results. Behav Nat Hum https://doi.org/10.1038/s41562-024-02046-9 (2024) xdoi:10.1038/s41562-024-02046-9.

40. Novikov, A. et al. AlphaEvolve: A coding agent for scientific and algorithmic discovery. Preprint at 10.48550/arXiv.2506.13131 (2025).

41. Romera-Paredes, B. et al. Mathematical discoveries from program search with large language models. Nature 625, 468–475 (2024).

42. Castro, P. S. et al. Discovering Symbolic Cognitive Models from Human and Animal Behavior. 2025.02.05.636732 Preprint at 10.1101/2025.02.05.636732 (2025).

43. Shojaee, P., Meidani, K., Gupta, S., Farimani, A. B. & Reddy, C. K. LLM-SR: Scientific Equation Discovery via Programming with Large Language Models. Preprint at 10.48550/arXiv.2404.18400 (2025).

44. Hammond, P. & Andrews, D. P. Orientation tuning of cells in areas 17 and 18 of the cat’s visual cortex. Exp Brain Res 31, 341–351 (1978).

45. Rose, D. & Blakemore, C. An analysis of orientation selectivity in the cat’s visual cortex. Exp Brain Res 20, 1–17 (1974).

46. Peyrache, A., Lacroix, M. M., Petersen, P. C. & Buzsáki, G. Internally organized mechanisms of the head direction sense. Nat Neurosci 18, 569– 575 (2015).

47. Andrews, D. P. Perception of contour orientation in the central fovea. I: short lines. Vision Res 7, 975–997 (1967).

48. Westheimer, G. & McKee, S. P. Spatial configurations for visual hyperacuity. Vision Res 17, 941–947 (1977).

49. Bradley, A., Skottun, B. C., Ohzawa, I., Sclar, G. & Freeman, R. D. Visual orientation and spatial frequency discrimination: a comparison of single neurons and behavior. J Neurophysiol 57, 755–772 (1987).

50. Rubin, D. B., Van Hooser, S. D. & Miller, K. D. The Stabilized Supralinear Network: A Unifying Circuit Motif Underlying Multi-Input Integration in Sensory Cortex. Neuron 85, 402– 417 (2015).

51. Ringach, D. L., Hawken, M. J. & Shapley, R. Dynamics of orientation tuning in macaque primary visual cortex. Nature 387, 281–284 (1997).

52. Wang, X., Lu, T., Snider, R. K. & Liang, L. Sustained firing in auditory cortex evoked by preferred stimuli. Nature 435, 341–346 (2005).

53. Stringer, C., Michaelos, M., Tsyboulski, D., Lindo, S. E. & Pachitariu, M. High-precision coding in visual cortex. Cell 184, 2767-2778.e15 (2021).

54. Stringer, C. et al. Spontaneous behaviors drive multidimensional, brainwide activity. Science 364, 255 (2019).

55. Goldey, G. J. et al. Removable cranial windows for long-term imaging in awake mice. Nat Protoc 9, 2515–2538 (2014).

56. Pachitariu, M. et al. Suite2p: beyond 10,000 neurons with standard two-photon microscopy. BioRxiv 061507, (2016).

57. Demas, J. et al. High-speed, cortex-wide volumetric recording of neuroactivity at cellular resolution using light beads microscopy. Nat Methods 18, 1103–1111 (2021).

58. Schmutz, V. et al. High-dimensional neuronal activity from low-dimensional latent dynamics: a solvable model. 2025.06.03.657632 Preprint at 10.1101/2025.06.03.657632 (2025).

## References

1. Tao, T. An Epsilon of Room, I: Real Analysis: Pages from Year Three of a Mathematical Blog. (American Mathematical Society, 2022).

2. Zygmund, A. Trigonometric Series. (Cambridge University Press, Cambridge, 2003). doi:10.1017/CBO9781316036587.

3. Erdélyi, A. Asymptotic Expansions. (Courier Corporation, 1956).

4. Rubin, D. B., Van Hooser, S. D. & Miller, K. D. The Stabilized Supralinear Network: A Unifying Circuit Motif Underlying Multi-Input Integration in Sensory Cortex. Neuron 85, 402–417 (2015).

5. Ringach, D. L., Hawken, M. J. & Shapley, R. Dynamics of orientation tuning in macaque primary visual cortex. Nature 387, 281–284 (1997).

6. Wang, X., Lu, T., Snider, R. K. & Liang, L. Sustained firing in auditory cortex evoked by preferred stimuli. Nature 435, 341–346 (2005).

7. Bartho, P., Curto, C., Luczak, A., Marguet, S. L. & Harris, K. D. Population coding of tone stimuli in auditory cortex: dynamic rate vector analysis. Eur J Neurosci 30, 1767–78 (2009).

